# The Immunoregulatory Architecture of the Adult Oral Cavity

**DOI:** 10.1101/2024.12.01.626279

**Authors:** Bruno F. Matuck, Khoa L. A. Huynh, Diana Pereira, Quinn T. Easter, XiuYu Zhang, Meik Kunz, Nikhil Kumar, Aditya Pratapa, Brittany T. Rupp, Ameer Ghodke, Alexander V. Predeus, Lili Szabó, Stefan Hartmann, Nadja Harnischfeger, Zohreh Khavandgar, Margaret Beach, Paola Perez, Benedikt Nilges, Maria M. Moreno, Kang I. Ko, Rohit Singh, Purushothama Rao Tata, Sarah A. Teichmann, Adam Kimple, Sarah Pringle, Kai Kretzschmar, Blake M. Warner, Inês Sequeira, Jinze Liu, Kevin M. Byrd

## Abstract

The immunoregulatory architecture of human oral tissues remains poorly defined despite their central role as barrier interfaces. We present the first integrated single-cell and dual-platform spatial-proteotranscriptomic atlas of oral tissues, profiling >250,000 single-cell transcriptomes and >4 million spatially-resolved cells across 13 niches. Using our AI-enabled AstroSuite (TACIT, Constellation, STARComm, hist2omics), we defined tissue cellular neighborhoods (TCNs) and multicellular interaction modules (MCIMs) in health, revealing peri-epithelial fibroblast-centered hubs enriched for effector cytokines. We harmonized eight fibroblast subtypes (universal, immune, peri-epithelial, peri-vascular, peri-neural, APC-like, stress-responsive, and myofibroblasts) with stress-responsive subtypes partitioning between mucosae (Type I) and glands (Type II). Spatial multiomics mapped receptor–ligand circuits and showed mucosal stress-responsive fibroblasts as immunoregulatory hubs. In chronic periodontitis, niche-aware integration of healthy and diseased datasets revealed rewiring of fibroblast phenotypes and ligand::receptor networks into interdigitated inflammatory and reparative niches. Disease neighborhoods exhibited fragmentation, expansion of MHC-I⁺, MHC-II⁺, and PD-L1⁺ fibroblasts, and predicted spatial engagement with T cells at ectopic lymphoid structures. Drug2Cell analysis highlighted druggable stromal::immune networks. Together, this proteotranscriptomic atlas positions fibroblasts as central architects of structural immunity in human oral tissues and establishes a scalable framework for precision targeting of stromal::immune ecosystems across other barrier organs in health and chronic disease.

## INTRODUCTION (729)

The oral cavity is both barrier and sentinel^1^, serving as a dynamic, immunologically rich interface where the body balances constant environmental exposure with robust internal defense mechanisms^2,3^. As a “window to the body”^4,5^, it contains multiple tissue types, including masticatory, lining and specialized oral mucosae, major and minor salivary glands, and teeth^6^. Despite regional differences in morphology^7^, mucosae share a core architecture of stratified squamous epithelium supported by keratinocytes and underlying stroma populated by fibroblasts, neurovascular units, muscle, and diverse immune cells^8^. Juxtaposed salivary glands, including parotid, submandibular, sublingual, and hundreds of minor salivary glands, are in contact with specific mucosal niches^9^. Despite this proximity, each oral and salivary niche is structurally and functionally distinct^6^, and certain diseases preferentially affect specific sites^10^. However, the cellular and molecular mechanisms underpinning niche-specific disease patterns across these distinct oral tissues remain poorly understood^11^.

Beyond their role as physical barriers, oral mucosae and glands exhibit sensory^12,13^ and emerging immunoregulatory functions^14^. Within these tissues, structural cells, such as keratinocytes, fibroblasts, and vascular endothelial cells dynamically engage with immune cells^15,16^. Long considered passive tissue support cells, structural cells are increasingly recognized as active regulators of immune function; this concept of structural immunity acknowledges their capacity to mediate immune responses, maintain homeostasis, and coordinate tissue regeneration^17,18^. Notably, the oral cavity is daily exposed to mechanical, microbial, and dietary stressors that may elicit sub-pathologic inflammation^19^, a form of low-grade, adaptive immune activation, known in other contexts as parainflammation^20,21^. This intermediary state exists between basal homeostasis and classical inflammation, and its molecular programs and spatial organization are not well characterized. Considering this, fibroblasts, which are strategically positioned across epithelial, perivascular, and interstitial zones, appear to be ideal candidates to mediate dynamic inflammatory states.

While recent efforts have mapped fibroblasts across organs and disease states^22–25^, the oral cavity remains largely underrepresented from such atlases. It remains unknown whether fibroblasts orchestrate conserved immune signaling strategies across distinct mucosae and glands, or whether niche-specific subtypes encode unique basal, para-inflammatory, pathologic, and/or immunoregulatory roles. Equally unclear is how this fibroblast-anchored architecture is remodeled in chronic oral disease states, which often feature niche-specific disruption of immune regulation within fibroblast-rich stromal compartments. To address these gaps, we constructed the first integrated single-cell and spatial proteotranscriptomic atlas of adult oral and craniofacial tissues. This resource includes single-cell RNA sequencing data from >250,000 cells from 70 samples representing 13 distinct tissue niches, including oral mucosae, salivary glands, and dental pulp, compiled using new and previously published datasets from the Human Cell Atlas Oral & Craniofacial Bionetwork^6^. We identified major structural and immune cell types, characterized their niche-specific heterogeneity, and profiled ligand–receptor expression patterns, revealing that fibroblasts emerged as among the most dynamic interactive cell types, suggesting a central role in orchestrating tissue-specific immune signaling.

To resolve these interactions *in situ*, we created two complementary spatial proteotranscriptomics atlases. First, we performed same-slide, spatial proteotranscriptomics using a 40-plex spatial-proteomics panel (Phenocycler-Fusion) after running a custom 300-plex spatial-transcriptomics panel (Xenium). Second, we applied a similar 40-plex spatial-proteomics panel and a similar custom 300-plex spatial-transcriptomics panel (MERSCOPE) to sequential tissue sections. Both atlases were designed to target known fibroblast-driven receptor–ligand pairs and applied to 21 unique human samples, capturing nearly 4 million cells across six distinct oral and craniofacial niches. To analyze these data, we applied our recently published spatial analysis suite, AstroSuite^26–28^, including 1) *hist2omics* (single-cell histological and/or multiomic data alignment), 2) *TACIT* (rapid, hierarchical cell annotation), 3) *Constellation* (integrated neighborhood identification), 4) *STARComm* (receptor-ligand network analyses), and 5) *Astrograph* (spatial data visualization).

Using these datasets, we resolved eight harmonized fibroblast subtypes across oral tissues, including universal, immune, peri-epithelial, peri-vascular, peri-neural, APC-like, stress-responsive, and myofibroblasts. Among these, stress-responsive fibroblasts were a prominent population in healthy tissues and further divided into two transcriptionally distinct groups: Type I, enriched in salivary glands, and Type II, enriched in mucosae. Spatial multi-omics revealed that fibroblasts structure tissue neighborhoods and coordinate receptor–ligand signaling, with mucosal stress-responsive fibroblasts emerging as potent immunoregulatory hubs. In chronic periodontitis, niche-aware integration showed rewiring of fibroblast states into interdigitated inflammatory and reparative niches, with expansion of MHC-I⁺, MHC-II⁺, and PD-L1⁺ fibroblasts near tertiary lymphoid structures. *Drug2Cell* analysis further identified context-specific therapeutic opportunities targeting these immune interfaces. Collectively, these findings position fibroblasts as central immune architects of the oral cavity and establish a scalable framework for precision targeting of stromal–immune ecosystems in health and chronic disease.

## RESULTS: (4616)

### An Integrated Single-cell Transcriptomics Atlas of Human Oral Tissues

The diversity of human cell-types and states remains incompletely defined and outside of the tooth and periodontium^18,29^, no integrated atlas of the human oral and craniofacial tissues exists. We compiled 11 publicly available single-cell RNA sequencing (scRNAseq) datasets, including samples from across oral mucosa^30–35^, salivary glands^31,35–38^, and dental pulp^32,39,40^ (Figure 1a,b), and generated three new scRNA-seq datasets from labial mucosa, soft palate mucosa, and parotid glands to expand coverage and representation of mucosal and glandular niches. We harmonized clinical and sample metadata for all the samples (Supplementary Table 1). We also generated spatial-proteomics and spatial-transcriptomics data using FFPE samples (Figure 1c) to create two complementary spatial multiomic atlases: (1) spatial-proteomics (Phenocycler-Fusion) with same-section spatial-transcriptomics (Xenium) and (2) spatial-proteomics with sequential-section spatial-transcriptomics (MERSCOPE), enabling single-cell–resolved integration of proteomic and transcriptomic measurements (Figure 1d)^26^.

**Figure 1.**
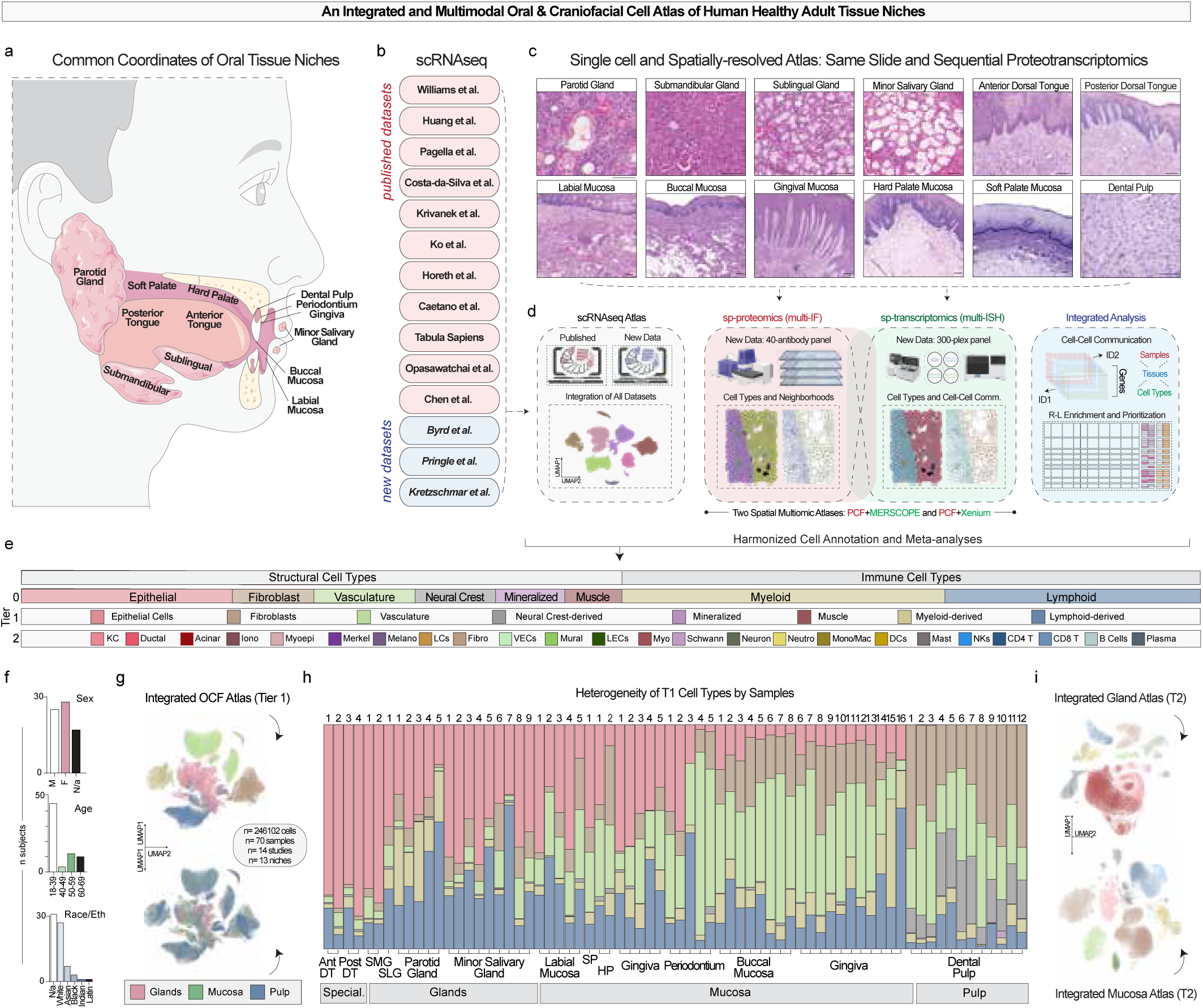
An Integrated and Multimodal Oral & Craniofacial Cell Atlas of Human Healthy Adult Tissues. (a) The common coordinate framework of oral and craniofacial macroniches. (b) Integration of publicly available scRNA-seq datasets (pink) and newly generated datasets (blue). (c) H&E images of niche-matched FFPE tissue curated for spatial analysis. (d) Overview of multimodal integration: the scRNA-seq atlas (gray); the multiplex immunofluorescence (spatial-proteomics, red); the multiplexed *in situ* hybridization atlas (spatial-transcriptomics, green). Integrated analyses (blue) combined. (e) Tiered annotation framework. (f) Demographics of donors in the scRNA-seq atlas. (g) UMAP plots of the integrated Tier 1 oral & craniofacial (OCF) atlas (n=246,102 cells from 70 samples, 14 studies, 13 niches), colored by cell-type (top) and tissue niche (bottom). Glands=pink, mucosa=green, pulp=blue. (h) Heterogeneity of Tier 1 cell-types across donor samples by niche. (i) Tier 2 subclustering of integrated gland and mucosal datasets, carried over from the integrated atlas in (g), with colors matched to Tier 1 categories in (e). Abbreviations: FFPE, formalin-fixed paraffin embedded, Iono, ionocytes, KC, keratinocyte, Myoepi, myoepithelial, Melano, melanocytes, LCs, Langerhans cells, Fibro, fibroblasts, VECs, vascular endothelial cells, Myo, myocytes, LECs, lymphatic endothelial cells, Neutro, neutrophils, Mono/mac, monocyte/macrophage, DCs, dendritic cells, NKs, natural killer cells, *T1*, Tier one. Scale bars: Glands: 100 µm; Mucosal 250 µm; pulp 50 µm (c).

To interpret the integrated scRNAseq integrated dataset and enable cross-niche comparisons, we next performed systematic cell-type annotation. We adopted a hierarchical classification approach to capture both broad lineage identities and finer niche-specific subtypes while maintaining consistency with established ontologies^41^. Following a tiered structure (Figure 1e), cells were classified into broad categories (Tier 0, T0) and refined into eight T1 types, which included structural (epithelial, fibroblast, vascular, neural crest-derived, muscle, mineralized) and immune (myeloid and lymphoid) populations. T1 annotations revealed consistent detection of most cell-types across samples.

The atlas encompasses three principal tissue groups: oral mucosae, salivary glands, and dental pulp, each with distinct cell-type compositions. Only healthy adult samples were included. The dataset skewed toward younger, female donors with limited racial and ethnic diversity (Figure 1f). After normalization and dimensionality reduction (Figure 1g), we benchmarked five batch-correction methods: (Harmony, Scanorama, BBKNN, FastMNN, and scVI) using scIB metrics for batch mixing, biological conservation, and cluster separation (Supplementary Figure 1a)^42^. Harmony achieved the best balance (score: 0.82) and was used for all subsequent analyses. After quality control (see *Methods*), the final atlas comprised 246,102 cells from 70-samples spanning 13-macroniches, publicly available via CELLxGENE (“Human Oral and Craniofacial Cell Atlas” / HCA), including subclustered atlases for major niches and cell-types. (Supplementary Figure 1b,c).

We computed cell-type signatures for each major cell-type (Supplementary Figure 1d). Epithelial cells were overrepresented in salivary gland and tongue samples, neural crest–derived populations were enriched in dental pulp, and vascular cells were enriched in mucosae (Figure 1h). Fibroblasts were present across samples, though their proportions varied by niche. UMAPs were then generated for all glands and mucosae (see *Data Availability*). T2 cell annotation across each tissue type increased the total number of identified cell-typest to 24, enhancing cellular resolution and revealing finer cell heterogeneity (Figure 1i), including some niche-specific epithelial types (glands: myoepithelial, ionocytes, acinar and ductal cells; mucosae: keratinocytes, melanocytes, Merkel and Langerhans cells). These data establish a foundational dataset of shared and niche-specific cell-types across oral tissues.

### Structural-Immune Communication Axes Across Oral Tissues

Having assembled the atlas, we first hypothesized that structural cells actively contribute to local immunoregulation through ligand::receptor (L::R) signaling and act as key initiators of immunomodulatory cues. To investigate this, we began by mapping the overall architecture of structural-to-immune cell communication. We first examined Tier 1 cell-type annotations in the version 1 atlas across tissues (Supplementary Figure 1e-g). The highest-ranked structural-to-immune interactions were dominated by extracellular matrix (ECM)-derived signals, such as *DCN*, *LUM*, *COL1A2/3A1*, *SPARC*, *PTN*, and *MFAP2* (Supplementary Figure 1h). We next compared Tier 1 cell-types between oral mucosa and salivary glands, which differ in function, to determine whether their structural::immune interaction profiles would also vary.

Adapting MultiNicheNet to eight glandular and mucosal niches (see: *Methods*), we modeled ligand::receptor signaling across samples and niches simultaneously^43^. We again identified a prominent fibrovascular signaling axis that varied between tissue types (Figure 2a,b; Supplementary Table 2). Across niches, fibroblasts engaged in extensive but diverse ECM signaling, underscoring their known roles as key architects of tissue organization. For example, in salivary glands, fibroblast::epithelial communication was anchored by *FGF2::FGFRL1* and *FGF10::FGFRL1*, pathways central to epithelial growth and ductal maintenance. Reciprocal Epithelial::fibroblast signals, including *EGF::EGFR* and *ANG::EGFR*, reinforced bidirectional signaling circuits. Fibroblast::vascular endothelial cell (VEC) interactions, such as *HBEGF::EGFR*, underscored fibrovascular crosstalk with potential roles in angiogenesis (Supplementary Figure 2a). In oral mucosae, fibroblast::epithelial signaling was dominated by *TNC::SDC1* and *COL3A1::ITGA1*. Fibroblast::fibroblast modules, including *TNC::ITGB1*. vascular::fibroblast (*JAM2::ITGB1*) and fibroblast::vascular (*VCAN::SELP*) interactions highlighted bidirectional activity that could be related to immune cell trafficking.

**Figure 2.**
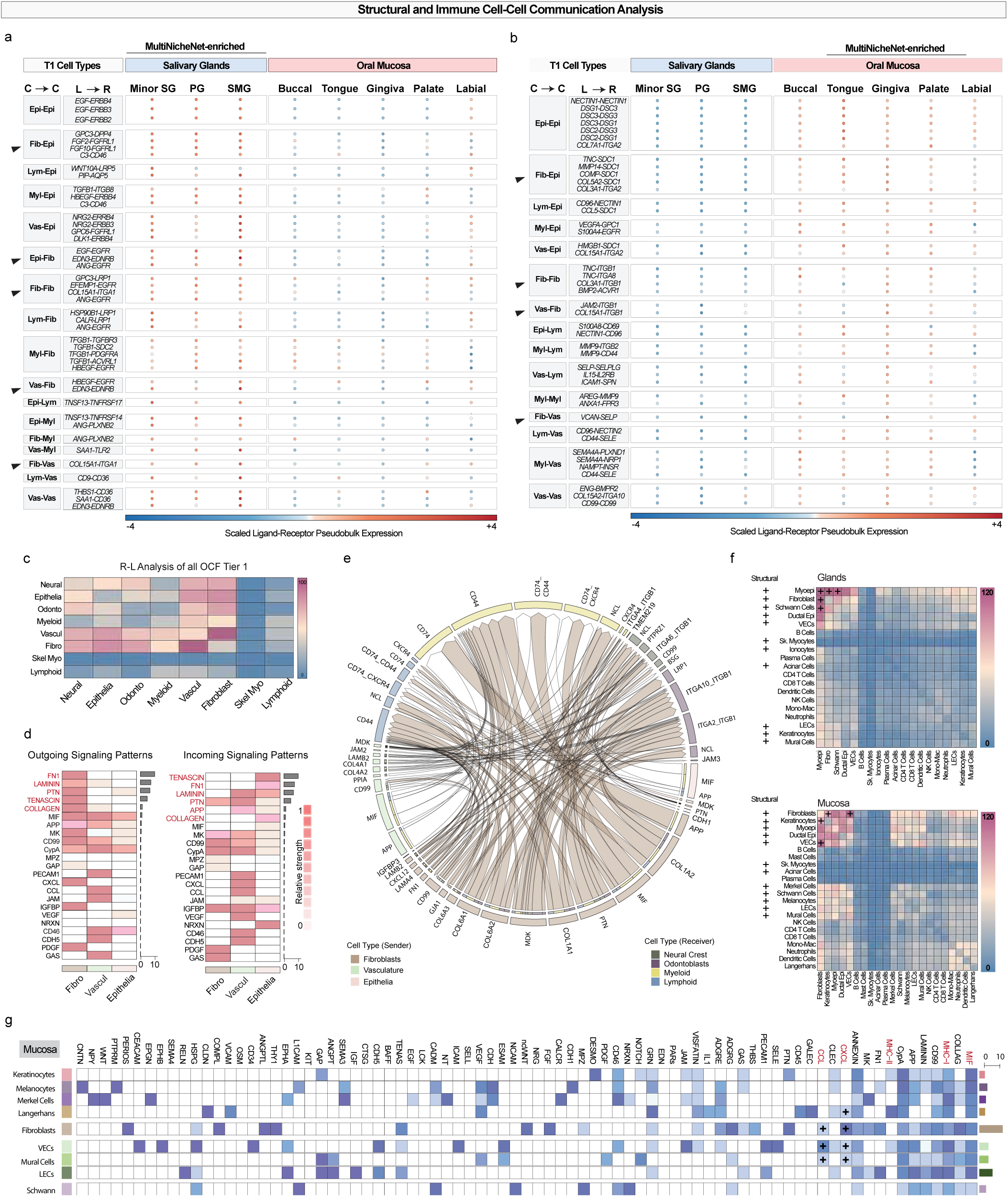
Structural and Immune Cell::Cell Communication Across Salivary Glands and Oral Mucosa. (a, b) MultiNicheNet-enriched ligand::receptor (L::R) analyses showing the top differentially regulated pairs between salivary glands and oral mucosae. (a) Communication enriched in glands relative to mucosa; (b) communication enriched in mucosa relative to glands. L::R interactions were inferred using CellPhoneDB and summarized as scaled pseudobulk expression values. C → C = cell-type sender::receiver relationship; L → R = ligand::receptor. Rows are grouped by structural and immune cell-types, with blue indicating lower and red indicating higher scaled expression. (c) Global L–R interaction heatmap across all T1 cell-types. (d) Outgoing (ligand) and incoming (receptor) signaling pathway patterns from CellChat analysis, highlighting enriched immune::structural communication signatures. (e) Chord diagram of the top 50-significant L–R interactions across all niches, illustrating directional communication between structural and immune cell-types. (f) Heatmaps showing total number of interactions in T2 structural-to-structural and structural-to-immune L–R signaling patterns across glands (top) and mucosa (bottom). (g) Pathway-level analysis of structural cell-communication, showing enrichment of CXCL, CCL, MHC, and MIF immune-related signaling modules across epithelial, fibroblast, vascular, and other structural populations. Abbreviations: SG, salivary gland; Minor SG, minor salivary gland; PG, parotid gland; SMG, submandibular gland; Epith, epithelial; Fib, fibroblast; Vas, vascular; Lym, lymphoid; My, myeloid; VECs, vascular endothelial cells; LECs, lymphatic endothelial cells.

Using CellPhoneDB, we quantified overall ligand::receptor activity, focusing on these key structural cell-types within the integrated atlas^44^. Fibroblasts and VECs displayed the highest levels of interaction potential, followed by epithelial and neural populations, with immune cells acting predominantly as receivers (Figure 2c; Supplementary Table 3). Pathway-level analyses of outgoing (ligand) and incoming (receptor) signals supported these trends (Figure 2d). Fibroblasts were especially enriched in IGFBP-and CXC-motif chemokine pathways, while vasculature utilized CC-motif chemokines. Using CellChat, we identified shared signaling circuits among epithelia, fibroblasts, and vasculature, often partitioning between ECM and immunomodulatory signaling (*Methods*; Figure 2e).

To capture structural::immune interaction diversity, we reanalyzed data with refined T2 cell-type resolution, revealing enriched networks in oral mucosae (Supplementary Figure 2b). L::R patterns across glands and mucosa again showed fibroblasts and VECs as major hubs, ranking top 5 in both (Figure 2f; Supplementary Table 3). In glands, fibroblasts signaled with epithelial cells, suggesting roles in architecture and homeostasis. In mucosa, fibroblasts interacted with keratinocytes and myeloid cells via chemokine pathways (Figure 2g), including CCL2::CCR2 (monocyte/DC recruitment), CCL5::CCR1 and CXCL12::CXCR4 (mast/NK interactions), and CCL19::CCR7 (lymphocyte homing). Fibroblasts were also predicted to engage MHC-I and CD99-mediated pathways (Supplementary Figure 2c), linking architecture to immunoregulation.

### Fibrovascular-anchored Neighborhoods Coordinate Divergent Immune States among Niches

We reasoned that the spatial organization of oral mucosa and salivary glands not only reflects tissue-specific structure but also actively shapes and sustains immune activity through coordinated interactions between structural and immune compartments. Informed by our T2 scRNAseq data, we generated a high-resolution spatial-proteomic atlas using a custom-designed panel of 40 oligo-conjugated antibodies applied to 6 healthy oral tissue types: 3 mucosal niches (tongue, gingiva, buccal) and three glandular (parotid, submandibular, minor; Figure 3a; Supplementary Figure 3a-c). Tissues were selected to have comparable demographics across groups (Supplementary Table 3).

**Figure 3.**
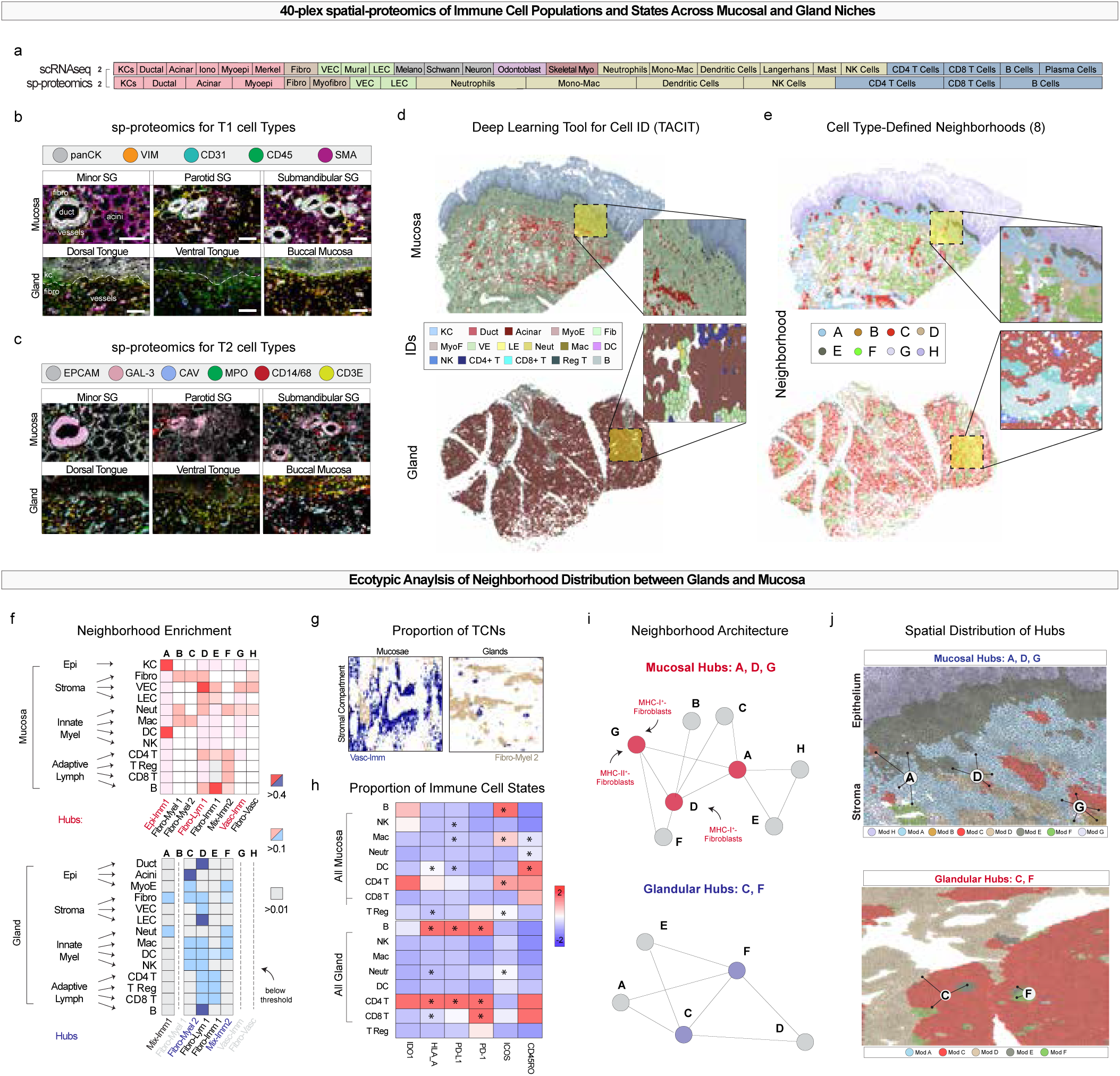
Spatial-proteomics Mapping of Immune and Stromal Neighborhoods Across Oral Mucosa and Salivary Glands. (a) Schematic of Tier 1 and Tier 2 cell-type classifications. (b–c) Representative multiplex immunofluorescence images showing Tier 1 (b) and Tier 2 (c) annotations in mucosa and glands. Cell-types were annotated using TACIT (AstroSuite; see Methods). (d) Deep-learning-based single-cell classification with TACIT, enabling full-tissue mapping of cell populations. (e) Tissue Cellular Neighborhoods (TCNs) identified after whole experiment integration by *Constellation*, assigning 8 distinct neighborhoods. (f) Neighborhood enrichment analysis showing epithelial, stromal, and immune composition within TCNs in mucosa (top) and glands (bottom). Neighborhood hubs, which are defined as TCNs with interactions to ≥2 other neighborhoods, are indicated. (g) Quantification of TCN proportions across mucosa and glands. (h) Immunophenotyping of distinct immune cells show divergent phenotypes by niche. (i) Network diagrams showing neighborhood architecture and hub connectivity, illustrating fibrovascular hubs in glands (C, F) and mucosa (A, D, G). (i) Spatial mapping of mucosal and glandular hubs, with annotated TCNs and surrounding neighborhood structures. Abbreviations: SG, salivary gland; VEC, vascular endothelial cell; TCN, tissue cellular neighborhood; Myofib, myofibroblast; Fibro, fibroblast; Mac, macrophage; DC, dendritic cell. Scale bars: 50 µm (b,c) (j). Abbreviations: TCN, tissue cellular neighborhood; VEC, vascular endothelial cell; Fib, fibroblast.

The antibody panel was validated across disease and health contexts using the Akoya Phenocycler-Fusion 2.0 platform, and Organ Mapping Antibody Panels (OMAPs) for these six tissues were included as part of the 9^th^ Release (v2.3) of the Human Reference Atlas^45^. Whole-slide images were segmented using Cellpose3 with H&E-guided, human-in-the-loop training (Supplementary Figure 3d)^46^. Initial results revealed marked differences in cellular composition and metacellular architecture between mucosal and glandular tissues, suggesting fundamentally distinct structural–immune landscapes (Figure 3b,c; Supplementary Figure 3e,f).

To enable such analyses at scale, we developed a modular and containerized spatial analysis toolkit: AstroSuite^26–28^, which includes: TACIT, a machine learning-based classifier for cell-types and states; Constellation, a framework for integrating and identifying Tissue Cellular Neighborhoods (TCNs); STARComm, a computational pipeline for mapping receptor–ligand signaling into multicellular immune modules (MCIMs) and organizing them into spatially resolved communication networks; and Astrograph, a tool for spatial data visualization (Figure 3d). Across both spatial-proteomics runs, including 36 total tissues, we identified ∼2,000,000 segmented cells that were used to annotate cell-types, states, and neighborhoods. As expected, mucosae displayed a higher proportion of innate immune cells (e.g., macrophages, *p*=0.012), while glands were enriched for adaptive cells (e.g., CD8⁺ T cells, *p*=0.006; Supplementary Figure 3g). Furthermore, using co-occurrence analysis, we found immune cells were significantly closer to structural cells in mucosae than in glands (Supplementary Figure 3h), indicating tighter immune– structural coupling at barrier surfaces.

To define TCNs, we scanned overlapping tissue regions and integrated patterns into consensus neighborhoods using Constellation (Figure 3f), identifying eight TCNs (A–H) across the dataset. Mucosae exhibited all eight, while glands harbored only five (A, C–F). Fibroblasts were enriched in four mucosal TCNs (B, C, D, H), with D and H also enriched in VECs, suggesting fibrovascular TCNs resembling “adventitial fibroblasts” from recent atlases. Innate immune cells were broadly distributed across TCNs, while adaptive immune cells preferentially localized to fibrovascular TCNs. Some TCNs were more abundant in mucosa than glands (Figure 3g), often at peri-epithelial zones, suggesting niches for innate immune surveillance and epithelial barrier maintenance. Divergence of TCNs and associated cell-types also showed divergent cell states (Figure 3h)..

We next analyzed spatial connectivity between TCNs to identify architectural network organization. We defined TCN “hubs” as neighborhoods connected to ≥3 others (Figure 3i). In mucosae, TCN-A, -D, and -G emerged as major hubs, representing mixed-immune, fibro-lymphoid, and vascular-immune neighborhoods, respectively. In glands, TCN-C and -F served as central hubs. While fibrovascular hubs were present in both tissues, those in mucosa were more spatially fragmented, reflecting potential para-inflammatory phenotypes related to structural-immune rewriting of the local niche (Figure 3j). Furthermore, we found higher expression of MHC-I^+^ (HLA-A^+^) fibroblasts in mucosal TCN hub D and MHC-I^+^ and MHC-II^+^ (HLA-DR^+^) fibroblasts in mucosal TCN hub G (Supplementary Figure 3i).

### scRNA-seq Resolves Fibroblast Diversity and Niche-Specific Programs Across Oral Tissues

Building on these spatially-defined fibrovascular hub phenotypes, we next sought to resolve the underlying fibroblast diversity and transcriptional states using scRNA-seq to determine whether specific subtypes align with the structural–immune architectures observed *in situ*. To achieve this, we first subclustered all fibroblasts from our harmonized scRNA-seq atlas, comprising >25,000 fibroblasts from 10 niches. Using literature-informed annotations, we resolved eight transcriptionally distinct subtypes (Figure 4a-c; Supplementary Figure 4a): Universal, Inflammatory, Perivascular, Myofibroblast, Peri-Epithelial, Peri-Neural, Stress-Responsive, and APC-like fibroblasts, each enriched in different niches with predicted unique functions^23,24,47–52^.

**Figure 4.**
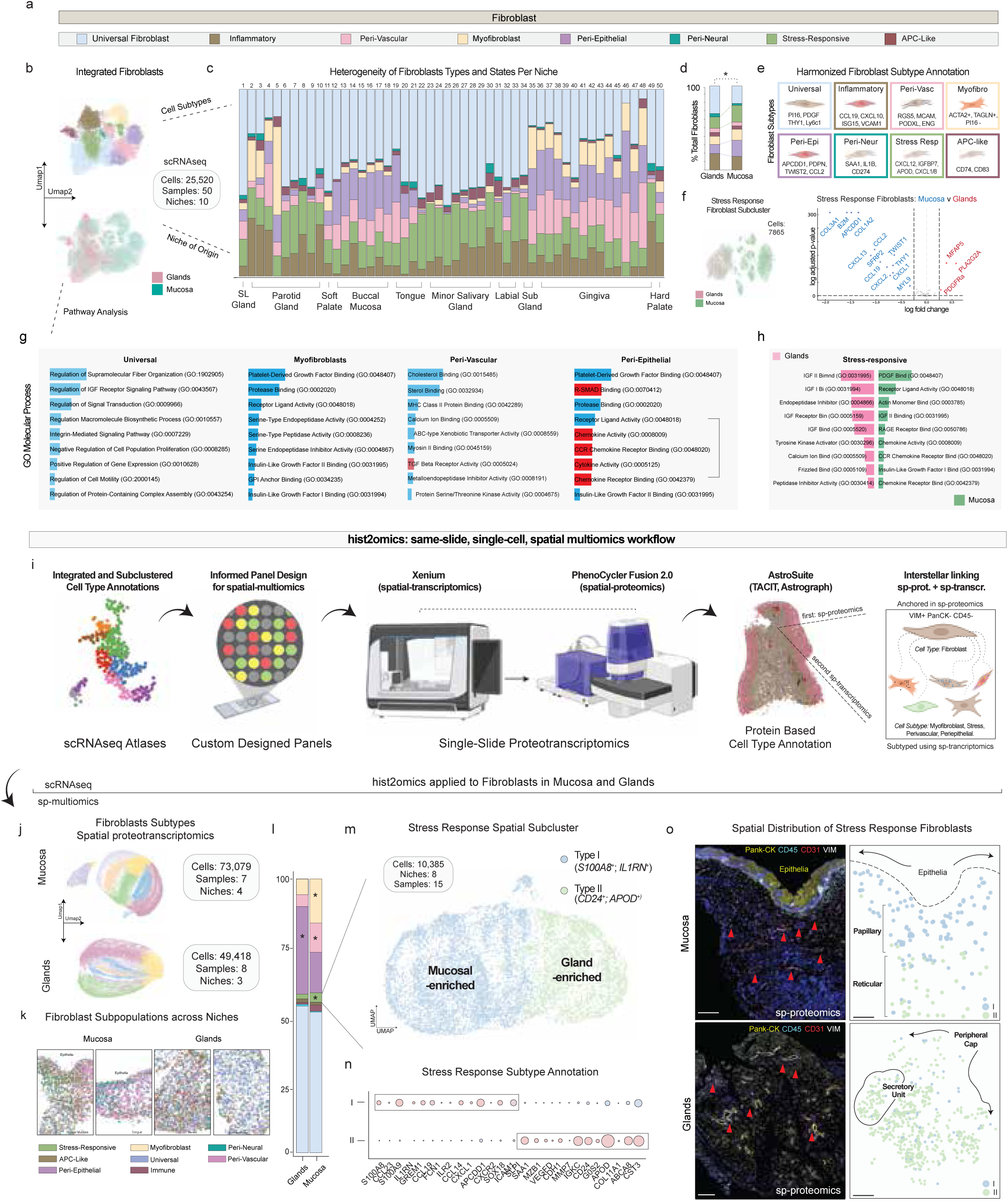
Fibroblast Diversity and Niche-Specific Polarization Across Oral Mucosa and Glands Revealed by scRNA-seq and Spatial Proteotranscriptomics. | (a) Fibroblast subtype color key, showing eight Tier 3 subtypes defined by integrated scRNA-seq. (b) UMAP projection of >25,000 fibroblasts from 50 samples and 10 oral niches, colored by subtype (top) or tissue of origin (bottom). (c) Proportional distribution of fibroblast subtypes per niche. (d) Overall subtype composition in glands versus mucosa; peri-epithelial and stress-responsive fibroblasts were enriched in mucosa. (e) Marker gene signatures used for fibroblast subtype annotation, highlighting distinct molecular programs. (f) Differential expression between mucosa and gland-enriched stress-responsive fibroblasts. (g–h) GO term enrichment analyses for universal, myofibroblast, peri-vascular, and peri-epithelial fibroblasts (g), and stress-responsive fibroblasts stratified by tissue. (h). Red color scale highlights immune pathways achieving significance, showing immune surveillance signatures in mucosa versus reparative, metabolism-linked programs in glands. (i) Hist2omics workflow: Linking spatial-transcriptomics and spatial-proteomics on the same slide. Cell annotation anchored by Vimentin⁺, PanCK⁻, CD31⁻, CD45⁻ cells for transcriptomic deeper subtype annotation. (j) UMAP projections of spatially annotated fibroblasts from mucosa (73,079 cells, 7 samples, 4 niches) and glands (49,418 cells, 8 samples, 3 niches). (k) Spatial maps of fibroblast subtype distribution within representative mucosal and glandular niches. (l) Quantification of subtype abundance across tissues, showing niche-specific enrichment patterns. (m) Stress-responsive fibroblast subclustering resolves two populations: Type I (mucosa-enriched) and Type II (gland-enriched). (n) Marker gene expression profiles distinguishing Type I (S100A8, IL1RN) and Type II (CD24, APOD) stress-responsive fibroblasts. (o) Spatial localization of Type I and Type II fibroblasts, showing peri-epithelial (papillary) dominance in mucosa and peri-epithelial plus deeper reticular positioning in glands *arrow:* Small Vessels. Scale Bar: 500 µm. Abbreviations: PCF, PhenoCycler-Fusion; VIM, Vimentin; PanCK, Pan-Cytokeratin; VEC, vascular endothelial cells; GO, Gene Ontology.

Proportional analysis demonstrated both shared and unique distributions by niche (chi-square; p<0.001; Figure 4d,e). Mucosal fibroblasts contained higher proportions of stress-responsive fibroblasts (*CXCL12⁺, IGFBP7⁺, APOD⁺, CXCL1⁺, CXCL8⁺*), whereas glands were enriched for universal (*P16⁺, PDGF⁺, THY1⁺*) and inflammatory fibroblasts (*CCL19⁺, CXCL10⁺, ISG15⁺, VCAM1⁺).* Additional subtypes included peri-epithelial (*APCDD1⁺, PDPN⁺, TWIST2⁺, CCL2⁺*), myofibroblasts (*ACTA2⁺, TAGLN⁺, PI16⁻*), APC-like fibroblasts (*CD74⁺*, *CD83⁺*), peri-neural fibroblasts (*SAA1⁺, IL1B⁺, CD274⁺*), and peri-vascular fibroblasts (*ACTA2^-^*, *RGS5⁺, MCAM⁺, PODXL⁺, ENG⁺*), each with molecular programs suggestive of specialized roles (Figure 4e, f). Stress-responsive fibroblasts comprised ∼25% of all fibroblasts, yet subclustering revealed divergent niche associations. Mucosa-enriched stress-responsive fibroblasts upregulated matrix remodeling genes (*COL1A2^+^, COL3A1^+^, MYL9^+^*), immune modulators (*CXCL1^+^, CXCL2^+^, CXCL13^+^, CCL2^+^, CCL19^+^*), and epithelial–mesenchymal regulators (*TWIST1^+^, SFRP2^+^, APCDD1 ^+^*).

By niche, we observed differences as well, highlighting the plasticity and dynamic potential of fibroblast populations across oral niches, supporting context-dependent functional specialization within shared structural archetypes. We were interested in the stress-responsive state within both mucosal and gland populations, as they were about 25% of the fibroblasts and upon subclustered analyses they displayed divergent clustering by niche (Figure 4f). Notably, mucosa-enriched stress-responsive fibroblasts displayed coordinated upregulation of matrix remodeling genes (*COL1A2, COL3A1, MYL9*), immune modulators (*CXCL1, CXCL2, CXCL13, CCL2, CCL19*), and epithelial–mesenchymal regulators (*TWIST1, SFRP2, APCDD1*), suggesting a hybrid state tuned for barrier maintenance, inflammatory signaling, and dynamic tissue remodeling as suggested by spatial-proteomics data (Figure 3). Consistent with our scRNAseq findings, this mucosal stress-response program may underpin heightened immune vigilance and epithelial resilience in barrier tissues, contrasting with the gland-predominant fibroblast states enriched for homeostatic or inflammatory roles.

Pathway enrichment analysis further underscored these niche-specific programs, with universal fibroblasts enriched for supramolecular fiber organization and IGF receptor signaling. Myofibroblasts were enriched for PDGF binding and protease activity, consistent with contractile and matrix-remodeling phenotypes that enable tissue reinforcement during repair. Peri-vascular fibroblasts displayed cholesterol/sterol binding and MHC class II activity, implicating them in lipid trafficking to vascular endothelium and potential perivascular antigen presentation. Peri-epithelial fibroblasts also exhibited PDGF binding together with pronounced chemokine and cytokine activity, positioning them as key modulators of epithelial proliferation and immune cell recruitment at barrier surfaces (Figure 4g).

Focusing on stress-responsive fibroblasts, mucosa-enriched populations showed preferential enrichment for PDGF binding, receptor–ligand activity, actin monomer binding, and RAGE/chemokine-receptor binding, all hallmarks of immune surveillance. In contrast, gland-predominant stress-responsive fibroblasts favored IGF-I/II binding, endopeptidase inhibition, tyrosine kinase activation, and calcium binding, suggesting a shift toward metabolically supported reparative programs that preserve secretory architecture and function under stress (Figure 4g, h). Taken together, these scRNA-seq findings resolve a fibroblast taxonomy with distinct niche-specific transcriptional programs and predicted functional polarizations, linking structural localization to specialized roles in ECM dynamics, immune modulation, and tissue-specific resilience. This data also suggests that niche-stress on fibroblasts can be intrinsic and distinct.

### Spatial Validation of Fibroblast Subtypes and Niche-Dependent Polarization

We next tested whether these fibroblast states were spatially organized and preserved *in situ*. To achieve this, we established and applied a new hist2omics framework to same-slide spatial proteotranscriptomic data, using a custom Xenium panel targeting scRNAseq-informed fibroblast subtype markers (Figure 4i). The proteotranscriptomic datasets were overlaid at single-cell resolution through image-based registration using their respective DAPI-stained nuclear images^26,53,54^. Fibroblast identity was anchored to the spatial-proteomics dataset by selecting Vimentin/(VIM)⁺, CD31⁻, CD45⁻, Pan-CK⁻ cells, thereby excluding endothelial, immune, and epithelial lineages. Segmentation masks generated from spatial-proteomics were transferred to the spatial-transcriptomics dataset, enabling deeper annotations restricted to the Vimentin⁺ population.

This protein-first anchoring was essential to harmonizing with the scRNAseq data, as spatial-transcriptomics annotation alone failed to capture a substantial fraction of confirmed fibroblasts seen by proteomics (Supplementary data 4a-c), often pulling in other cell-types not intended to be studied here. Once filtered, VIM+, CD31⁻, CD45⁻, Pan-CK⁻ cells displayed similar signatures to scRNAseq that were overall consistent by niche (Supplementary Figure 4d,e). As expected, the effector cytokine expression was highest in our stress-responsive and inflammatory fibroblasts states as expected (Supplementary Figure 4f). Considering glands and mucosae, we found that the 8 fibroblast types were distributed logically between epithelial and stromal compartments (Supplementary Figure 4g). Ultimately, ∼122,000 fibroblasts were identified robust but exhibited niche-specific heterogeneity (Figure 4j). For example, mucosal peri-epithelial zones were enriched for peri-epithelial, inflammatory, stress-responsive, and myofibroblast populations, often forming spatial gradients toward epithelial borders. Glandular fibroblasts favored universal and immune-like phenotypes with more dispersed spatial patterns (Figure 4k). Analyses showed expansion of stress-responsive fibroblasts and myofibroblasts, modest enrichment of immune and peri-vascular fibroblasts in mucosa, and more peri-epithelial fibroblasts in glands, reflecting acinar and ductal dominance (Figure 4l).

We again focused on our newly defined and divergent stress-responsive fibroblasts, which represented approximately 10% of total fibroblasts in spatial data. Unsupervised subclustering distinctly resolved our stress responsive fibroblasts into two stress-state populations, now termed Type I-and Type II-stress responsive fibroblasts, with divergent niche distributions (Figure 4m). Type I-fibroblasts, enriched in mucosa as seen in scRNAseq data, expressed inflammatory surveillance, epithelial mesenchymal plasticity, and matrix remodeling genes (*S100A8/9, CCL19, IL1RN, CXCL1, GREM1, CXCR2, ICAM1*; Figure 4n). Type II-fibroblasts, predominant in glands, expressed repair, immune modulatory, and lipid metabolism genes (*CST3, ABCA8, COL11A1, APOD, CD24, IGKC, MMP7, VEGFD, SAA1, SLP1*). Spatial localization confirmed these niche-specific programs as Type I-fibroblasts localized primarily to peri-epithelial (papillary) layers of oral mucosae lamina propria, whereas Type II-fibroblasts were enriched in deeper reticular mucosal layers of lamina propria and peri-epithelial regions of glandular tissue (Figure 4o). These single-cell and spatial analyses show that fibroblast subtypes found by scRNA-seq are preserved in tissues and display niche-specific locations and functions.

### Spatial Proteotranscriptomics Predicts Fibroblast-Driven Interaction Hubs

Having identified fibroblast subtypes and zones of immune activation, we sought to determine whether fibroblasts were embedded within reproducible, spatially organized networks of cell::cell communication that could functionally define their niche roles. Inspired by frameworks such as tissue cellular neighborhoods^55^ and recently proposed spatial ecotypes^56^, we hypothesized that receptor-ligand interactions in oral tissues might also follow patterned architecture, forming clusters of interacting cells defined not just by type or location, but by their spatially coordinated and recurrent cell-cell communication activity. To answer this, we applied our STARComm framework to find L::R neighborhoods, what we define as multicellular interaction modules (MCIMs), to two complementary spatial-transcriptomics datasets in parallel (Figure 5a).

**Figure 5.**
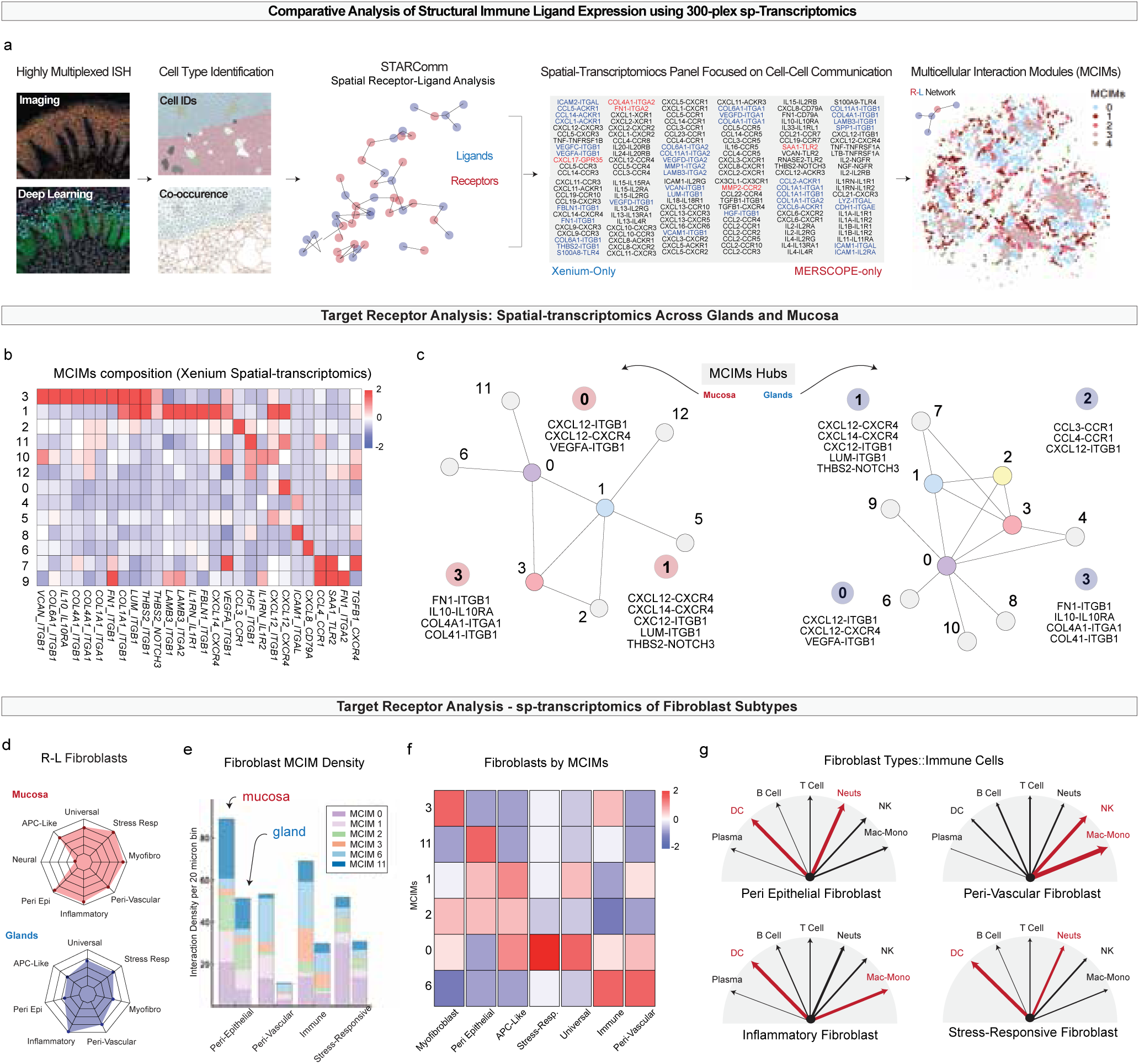
Spatial proteotranscriptomics reveals fibroblast-centered multicellular interaction modules (MCIMs) across oral mucosae and salivary glands. (a) Overview of the STARComm (SpaTiAl Receptor–Ligand Analysis for Cell–Cell Communication) workflow applied to Xenium (∼125 curated R–L pairs) and MERSCOPE (∼100 curated R–L pairs) datasets. (b) Heatmap showing MCIM composition from Xenium-derived spatial-transcriptomics across mucosal and glandular tissues. Rows represent individual MCIMs; columns list representative receptor–ligand pairs. (c) Network plots of MCIM hubs by tissue type, highlighting mucosa-enriched (red) and gland-enriched (blue) modules. (d) Radar plots showing receptor–ligand expression profiles of fibroblast subtypes in mucosa (top) and glands (bottom). Arrow thickness represents the CellChat communication score (range 0–1), red arrows denoting interactions with p < 0.05. (e) Distribution of fibroblast MCIM density by subtype and niche, showing the relative contribution of each fibroblast population to MCIM clusters. (f) Heatmap of fibroblast subtype participation across MCIMs, illustrating niche-specific module engagement (e.g., peri-epithelial fibroblasts with MCIM 11, myofibroblasts with MCIM 3). (g) Radar plots showing fibroblast subtype–immune cell interaction biases inferred from STARComm. Peri-epithelial and stress-responsive fibroblasts preferentially interact with DCs and neutrophils, whereas inflammatory and peri-vascular fibroblasts target monocyte/macrophage, DC, and NK cells.

In the first dataset, we used spatial-transcriptomics (MERSCOPE) with a 300-plex panel (∼100 curated L–R pairs, 20% of content) covering mucosae and glands (Supplementary Table 4), piloting STARComm with a single-modality assay. In the second, we applied hist2omics, a same-slide dual-modality workflow linking Xenium (spatial-transcriptomics) with Phenocycler-Fusion (spatial-proteomics) for protein-anchored annotation and high-plex RNA detection. The Xenium panel (∼125 curated R–L pairs, Supplementary Table 5) resolved fibroblast subtypes, with ∼25% fibroblast-relevant pairs absent from MERSCOPE. Figure 5a shows Xenium-only (blue) and MERSCOPE-only (red) pairs and illustrates how STARComm integrates in situ hybridization, TACIT-based annotation, spatial co-occurrence, and constrained R–L proximity analysis to cluster MCIMs.

MERSCOPE-derived STARComm analysis revealed 17 MCIMs across oral mucosae and glands (Supplementary Figure 5a). Mucosae supported 15/17 modules (missing Modules 2 and 13), while glands contained only 10/17, with some modules (e.g., Module 14) uniquely present in minor salivary glands. Several MCIMs were dominated by fibroblast::immune or fibroblast::vascular interactions. Modules 3 and 7, containing *CXCL12::CXCR4*, *CXCL16::CXCR6*, and *CCL14::CCR1* pairs, were highly represented in mucosa and reflected stress-responsive fibroblast (Type I) crosstalk with immune partners. Module 5, enriched in *IL10::IL10RA* and *EGF::EGFR*, linked fibroblasts with vascular and epithelial compartments. Fibroblasts were among the most highly signaling populations using a 50µm radius, with mucosal fibroblast-dominated MCIMs enriched for immune surveillance and barrier-maintenance pathways. We found niche-specific enrichment of MCIMs (Supplementary Figure 5b). Across all modules, clustering of R–L pairs (Supplementary Figure 5c) revealed distinct co-associations, suggesting recurrent signaling programs that span multiple MCIMs. For example*, CXCL12::CXCR4* clustered with several chemokine–chemokine receptor axes (e.g., *CCL17::CCR4, CXCL16::CXCR6*), consistent with coordinated lymphocyte recruitment from vascular networks.

To validate fibroblast-centered MCIMs considering our newly defined cell-types and states, we applied STARComm to the proteotranscriptomics dataset (Figure 4). We identified shared and distinct MCIMs across oral mucosae and glands. Across the 13 Xenium-derived MCIMs (Figure 5b), interaction signatures were dominated by two major themes similar to previous L–R analyses: (1) ECM–integrin signaling, with repeated enrichment of collagen (*COL1A1, COL4A1, COL6A1*), laminin (*LAMA5, LAMB3*), and fibronectin (*FN1*) binding to *ITGB1/ITGA1/ITGA2*, suggesting coordinated stromal–structural anchoring across both niches; and (2) inflammatory chemokine and cytokine axes, including *CXCL12::CXCR4, CXCL8::CD79A*, and *CCL4::CCR1* (Figure 5c). The differing number of identified MCIMs (17 in MERSCOPE; 13 in Xenium) reflects the non-overlapping L::R pair used in each dataset, driving finer module resolution and greater total module counts in MERSCOPE.

Mucosal tissues showed a broader distribution of modules (MCIMs 0, 1, 3, 6, 11, and 12), whereas glands were enriched for a distinct subset (MCIMs 0, 1, 2, 3, and 4; Supplemental Figure 5d,e). Mucosal MCIMs 1, 11, and 12 showed the highest spatial density, defined by fibroblast-mediated interactions such as: *HGF::ITGB1*, *COL4A1::ITGA1*, and *CXCL12::CXCR4*. Gland-associated MCIMs featured a narrower L::R repertoire. MCIM 0 was essentially driven by *CXCL12::CXCR4*, while MCIMs 2 and 4 centered around *CCL3::CCR1*, *ICAM1::ITGAL*, and *VEGFA::ITGB1*. These differences were consistent with prior observations of tissue cellular neighborhoods, with mucosa and glands maintaining overlapping yet distinct MCIM hubs (e.g., mucosal hubs: MCIMs 0, 1, 3; glandular hubs: MCIMs 0-3). Across both technologies, *CXCL12::CXCR4* emerged as the most conserved and spatially recurrent axis, linking fibroblast-rich niches with immune and vascular guidance across tissue types.

Same-slide proteotranscriptomics linked ligand transcripts to protein-anchored fibroblasts (Figure 4). Overall, ligands were higher in mucosal than glandular fibroblasts, with inflammatory states highest in both niches (Figure 5d). In mucosa, Type I stress-responsive, myofibroblast, and peri-epithelial subsets were also ligand-rich, unlike glandular fibroblasts. Mucosal peri-neural fibroblasts showed low L::R activity, likely due to panel limits. In glands, inflammatory fibroblasts dominated, with low counts in Type II stress-responsive cells.

Across the major signaling modules (0, 1, 2, 3, 6, and 11), these four shared (Peri-epithelial, Peri-Vascular, Immune, and Stress-Responsive) higher enrichments of module 0 (*CXCL12::CXCR4*) and module 6 (*CXCL8::CD79A*) (Figure 5e). When assessing fibroblast subtypes for confirmed L-R pair signatures (Figure 5f), myofibroblasts were enriched for module 3 (ECM ligands across COL, FN, VCAN), peri-epithelial fibroblasts for module 11 (*HGF::ITGB1*), and APC-like fibroblasts for *LAMININ, IL1RN*, and *CXCL12*. Stress-responsive and universal fibroblasts shared signatures with inflammatory and peri-vascular fibroblasts (module 6). Biologically, inflammatory and peri-vascular fibroblasts preferentially signal to monocytes/macrophages and NK cells. The highest L-R linked axis for T cells was centered on peri-vascular (Figure 5g). Peri-epithelial and Type-I/II stress-responsive fibroblasts skewed toward an innate immune axis (preferential dendritic cell and neutrophil interactions). By linking fibroblast subtype-resolved hist2omics with STARComm, we mapped local signaling partners for nearly all fibroblast subtypes across innate and adaptive immune populations, suggesting that newly described stress responsive fibroblasts are supportive of innate immune residency within diverse oral tissue niches.

### Fibroblast-Led Immunoregulation and Spatial Network Remodeling in Chronic Periodontitis

We next applied this approach to interrogate fibroblast spatial architecture and immunoregulatory function in chronic inflammatory disease. Furthermore, we wanted to extend our STARComm-based framework into disease contexts. We again focused on fibroblast activity *in situ*, reanalyzing scRNA-seq datasets from our Version 1 human periodontitis scRNAseq atlas^18^; we also expanded our atlas to generate a multimodal map of health and disease in this niche using the same-slide multimodal (Figure 6a; Supplementary Figure 6,7a). To preserve spatial fidelity, tissue sections were consistently oriented along the tooth–epithelial interface. We hypothesized that chronic inflammatory pressure drives fibroblasts toward spatially-anchored immunomodulatory phenotypes that engage and regulate innate immune and lymphocyte-rich immune compartments.

**Figure 6.**
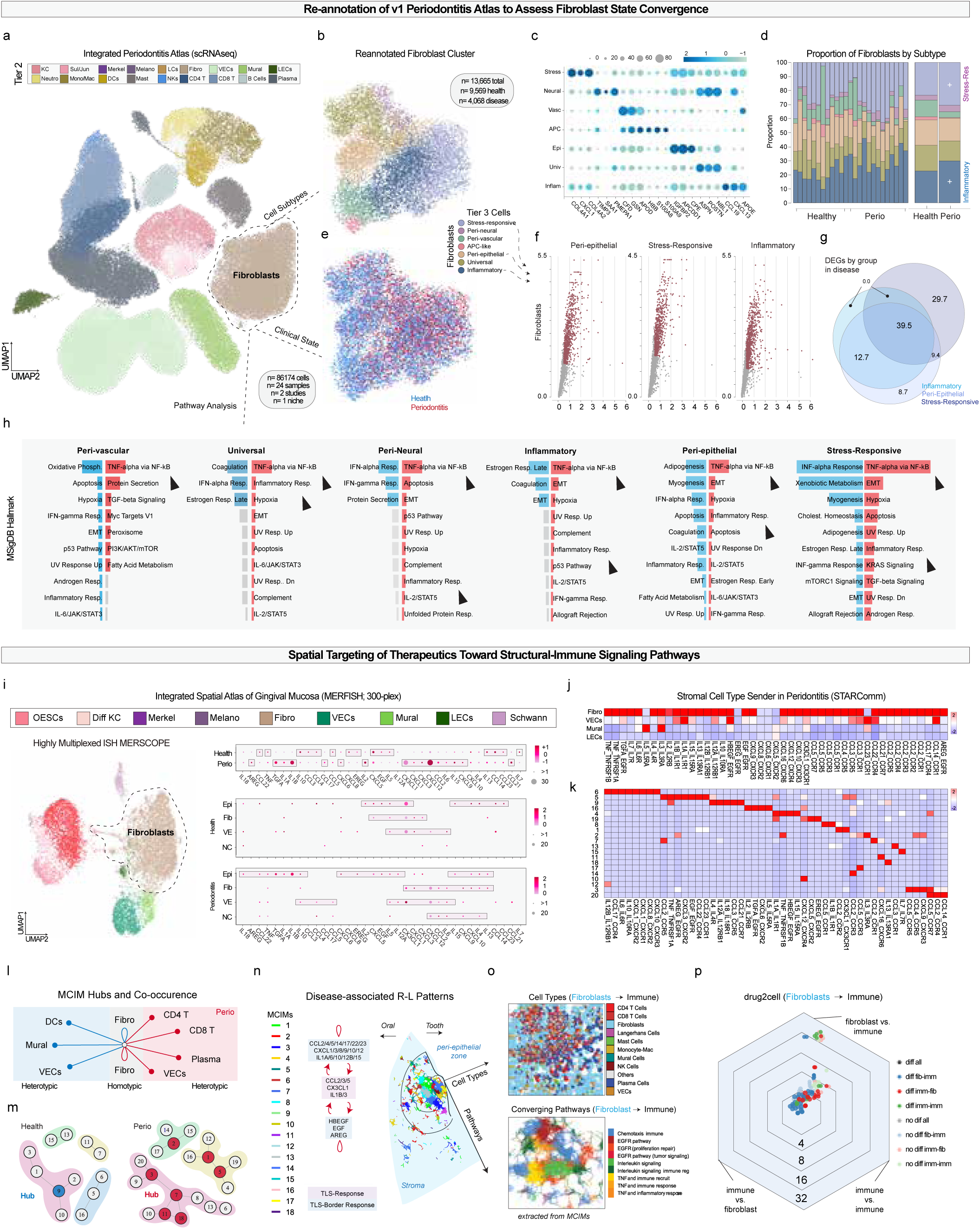
Fibroblast-Led Immunoregulation and Spatial Network Remodeling in Chronic Periodontitis. (a) Integrated periodontitis atlas (scRNA-seq) re-annotated using updated Tier 3 fibroblast state taxonomy. (b) UMAP of re-clustered fibroblasts subtypes (n = 13,665 cells) from healthy and periodontitis gingiva. (c) Dot plot of fibroblast subtype marker expression. (d) Proportion of fibroblasts by subtype in healthy vs. periodontitis samples, showing expansion of stress-responsive and inflammatory fibroblasts. (e) Tier 3 cell distribution across health and disease. (f) Volcano-plots showing disease-associated differential gene expression (DEGs) for peri-epithelial, stress-responsive, and inflammatory fibroblasts. (g) Venn diagram of shared and unique DEGs among fibroblast subtypes in disease. (h) MSigDB pathway enrichment in each fibroblast subtype, highlighting TNF-α, NF-κB, and TGF-β pathway activation in stress-responsive fibroblasts. Gray color indicates no significance achieved and term removed. (i) Integrated spatial atlas of gingival mucosa (MERSCOPE 300-plex) with fibroblast-focused ligand expression maps. (j) STARComm analysis of fibroblast-to-immune ligand–receptor interactions, showing expanded chemokine, interleukin, and EGFR-associated signaling in periodontitis. (k) Heatmap of fibroblast subtype–specific ligand–receptor interactions in health and disease. (l) MCIM hubs and co-occurrence networks in health vs. periodontitis, with disease-specific hubs enriched for *CCL2::CCR2, CX3CL1::CX3CR1, AREG::EGFR, TNF::TNFRSF1B,* and *IL3::IL3RA*. (m) MCIM modules in health vs. periodontitis, showing increase to 20 modules and fibroblast dominance in both homotypic and heterotypic interactions. (n) Spatial mapping of disease-associated MCIMs, highlighting TLS-enriched modules (1, 7, 8, 13, 14, 17) dominated by fibroblast chemokine and interleukin signaling, and surrounding modules (2–6, 20) enriched for cytokine and EGFR ligand pathways. (o) TLS-associated fibroblast programs showing combined immune recruitment, inflammation, and wound-repair activity within peri-epithelial zones. (p) Spatial *Drug2Cell* analysis of TLS-enriched fibroblast–immune ligand–receptor pairs. Abbreviations: APC-like, antigen-presenting cell-like fibroblasts; EGFR, epidermal growth factor receptor; IL, interleukin; MCIM, multicellular interaction module; MSigDB, Molecular Signatures Database; NF-κB, nuclear factor kappa-light-chain-enhancer of activated B cells; TGF-β, transforming growth factor beta; TLS, tertiary lymphoid structure

Applying our updated fibroblast annotation schema, we re-clustered fibroblasts across 24 samples (n=13,665 cells), recovering the same T3 subpopulations: stress-responsive, inflammatory, peri-epithelial, peri-neural, peri-vascular, antigen-presenting cell–like (APC-like), and universal fibroblasts (Figure 6b,c). In periodontitis, the proportions of stress-responsive and inflammatory fibroblasts appeared to expand, while universal, APC-like, and peri-neural populations contract (Figure 6d). When comparing genes upregulated in periodontitis compared to controls, most differences were found in inflammatory and stress-responsive clusters (Figure 6e,f). Peri-epithelial fibroblasts, which are adjacent to disease-originating niche in periodontitis stage III/grade B, shared substantial transcriptional overlap with these pro-inflammatory clusters, suggesting a plastic injury-response program near site of disease origination (Figure 6g). MSigDB analysis identified strong TNF-α, NF-κB, and TGF-β pathway activation in Type I-stress-responsive fibroblasts, the highest among subpopulations (Figure 6h), pointing to an underappreciated role for these new cell states in immunoregulation.

To spatially contextualize this fibroblast ligand expression (Supplementary Figure 7c,d), our MERSCOPE data was reanalyzed with a focus on structural::immune ligands (Figure 6i). We mapped fibroblast activity along the lesion interface, Importantly, in periodontitis, fibroblasts exhibited upregulation of pro-inflammatory mediators such as *TNF, IL1B, CXCL8*, and *AREG*, with fibroblasts showing the most pronounced signaling increases compared to other structural cel-types (LECs, mural cells, VECs; Figure 6j). L::R analyses revealed a marked expansion of fibroblast-directed immune signaling across chemokine, interleukin, and *EGFR*-associated modules (Figure 6k).

Fibroblasts remained the highest-predicted interaction partners, both homotypically (fibroblast::fibroblast) and heterotypically with immune cells (fibroblast::immune; Supplementary Figure 7e; Figure 6l). Considering tissue-wide rewiring in disease, the number of MCIMs increased from 13 in healthy mucosa to 20 in disease (Figure 6m). Disease-specific MCIMs (e.g., 3 and 7) showed expanded CD4⁺ and CD8⁺ T cell interactions. Disease hub modules were enriched for *CCL2::CCR2, CX3CL1::CX3CR1, AREG::EGFR, TNF::TNFRSF1B,* and *IL3::IL3RA*, many of which were fibroblast-specific. While individual fibroblast::immune axes have been described in other chronic inflammatory conditions, no prior periodontitis studies have mapped such disease-specific MCIMs linking fibroblasts to CD4⁺/CD8⁺ T cells within a spatially resolved multicellular network.

Spatial module analysis in and around tertiary lymphoid structures (TLS) revealed fibroblast– immune MCIMs concentrated both within and surrounding these immune aggregates (Figure 6n). TLS-enriched modules (1, 7, 8, 13, 14, 17) were dominated by fibroblast *CCL2, CCL5, CX3CL1,* and *IL1B* signaling, whereas surrounding modules (2–6, 20) featured broader cytokine and EGFR ligand signaling (*IL6, TNF, CXCL9/10/12, AREG, HBEGF*). TLS borders exhibited a patchwork of fibroblast programs that simultaneously recruited immune cells, promoted inflammation, and engaged wound-repair pathways, particularly within peri-epithelial zones (Figure 6o).

To identify therapeutic entry points, we filtered fibroblast–immune L-R pairs enriched in TLS zones and applied our spatial *Drug2Cell* workflow, revealing that nearly all were targetable with existing compounds (Figure 6p; Supplementary Figure 7f) Top nominated axes included GM-CSF/CSF2 (Sargramostim), IL-6 (Satralizumab), and IL-2RA/CD25 (Basiliximab), alongside T-cell–directed agents such as muromonab-CD3, consistent with fibroblast-linked immunoregulation in TLS-associated MCIMs (Supplementary Figure 7g). As *Drug2Cell* only nominates pharmacologically tractable axes by linking expressed targets to known drugs, without distinguishing agonist from antagonist strategies, the biological interpretation must consider context; for example, GM-CSF agonism could exacerbate chronic inflammation in gingiva, whereas antagonism of the same axis could have the opposite effect^57^. Mapping predicted drug sites of action back to tissue confirmed spatially restricted vulnerabilities within fibrovascular hubs.

We then examined our spatial-proteomics data in the context of fibroblast phenotypes to examine changes in TCNs, checkpoint ligand expression, and stromal–immune co-localization in periodontitis. All TCNs identified in health were retained in disease, but their spatial arrangement and cellular composition were markedly altered (Figure 7a,b). Additionally, a new VEC/NK cell-enriched TCN was added (TCN-B). The fibrovascular corridor, a structural region enriched in fibroblasts, endothelial cells, and immune populations, fragmented into smaller, spatially discrete TCNs. These localized hubs were found both in peri-epithelial regions adjacent to microbial insult and in peri-osteal zones near the alveolar bone. By immunophenotyping the TCNs, we found significantly upregulated BCL2, CD107A, PD-L1 and ICOS in TCN-G, a known fibrovascular hub (Figure 7c,d). This supports direct fibro-immune interactions were possible, and emergence of this TCN may serve as a biomarker of disease activity.

**Figure 7:**
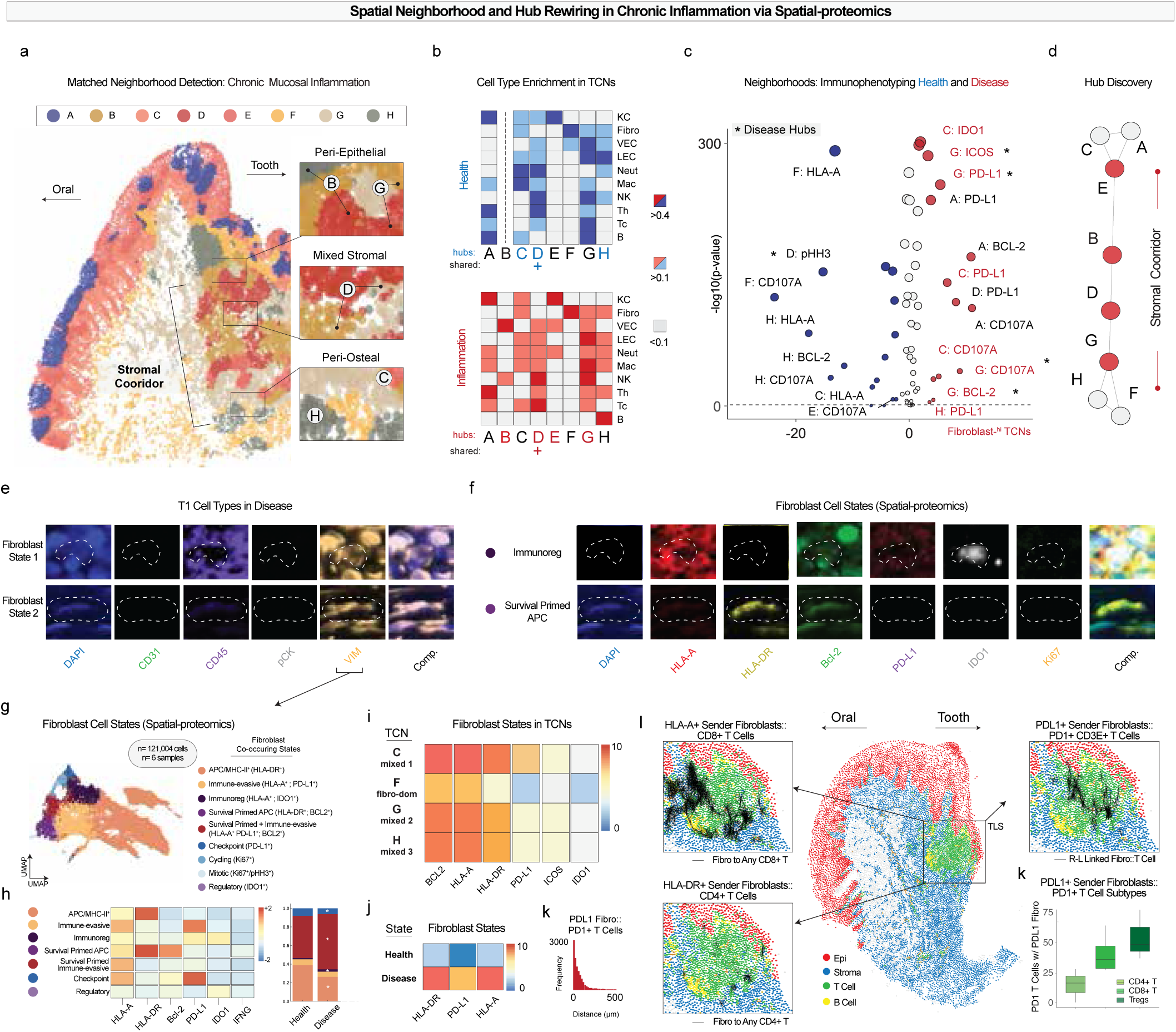
Spatial Neighborhood and Hub Rewiring in Chronic Inflammation via Spatial-proteomics. (a) Tissue-wide spatial mapping of matched tissue cellular neighborhoods (TCNs) in chronically inflamed oral mucosa reveals spatial reorganization of stromal–immune architecture. Neighborhood (e.g., peri-epithelial [B], peri-osteal [C]) and fragmentation of the stromal corridor are noted. (b) Cell-type enrichment heatmaps across TCNs in health (top) and disease (bottom) showing altered distribution of immune (T, B, NK, Myeloid) and stromal (Fib, VEC, LEC) cell-types. (c) Volcano-plot of immunophenotypic markers across TCNs showing upregulation of immune checkpoint and survival markers (e.g., PD-L1, IDO1, BCL2) in specific fibroblast-enriched neighborhoods. (d) Network diagram illustrating disease-driven remodeling of fibrovascular hub connectivity among TCNs, with G emerging as a prominent immune-enriched hub. (e,f) Multiplex immunofluorescence focused on fibroblasts (e) and their states (f) reveals diversity of expression, expressing immunoregulatory molecules. (g) Spatial map of fibroblast cell states using spatial-proteomics across inflamed oral mucosa, revealing distribution of 9 immunoregulatory states. (h) Heatmap of marker combinations across spatially defined fibroblast states and their frequency across health and disease. Stacked bar graph showing proportion Survival Primed + Immune-evasive (HLA-A+ PD-L1+; BCL2+) as well as Immunoregulatory (HLA-A+ ; IDO1+) expands states in disease. (i) Distribution of fibroblast states across disease-associated TCNs showing fibroblast heterogeneity in inflamed hubs. (j) Overall, independent of TCNs, all three MHCI/II and PDL1 molecules increase in disease. (k) PD-L1 (fibroblasts)::PD-1 (CD3 T Cells) are frequently collocated *in situ*. (l) Spatial co-localization of fibroblast–T cell checkpoint interactions. Left top and bottom: fibroblast sender cells (red) and T cell targets (blue) for HLA-A/CD8 and HLA-DR/CD4 axes; right top is and image of PD-L1 (fibroblasts)::PD-1 (CD3 T Cells) predictions. (m) Histogram showing highest PD-L1::PD-1 pairing in regulatory and cytotoxic CD3⁺ T cells in this data.

Interrogation of the scRNA-seq dataset for an expanded immunoregulatory gene panel (Supplementary Data 7h) suggested that fibroblasts could express MHC class I/II molecules and multiple immune checkpoint molecules. In APC-like fibroblasts, low-level but co-occurring expression of MHC molecules, PD-L1, and IDO1 was consistent with a capacity for both antigen presentation and immune evasion. To directly assess these features *in situ*, we subclustered fibroblasts from the spatial-proteomics dataset and profiled them for immunophenotypes via multiplex imaging (HLA-A, HLA-DR, BCL2, PD-L1, IDO1, Ki67) (Figure 7e,f). This analysis identified nine major fibroblast states defined by immunoregulatory protein expression and/or proliferative activity, including APC/MHC-II⁺ (HLA-DR⁺), immune-evasive (HLA-A⁺, PD-L1⁺), immunoregulatory (HLA-A⁺, IDO1⁺), survival-primed APC (HLA-DR⁺, BCL2⁺), survival-primed + immune-evasive (HLA-A⁺, PD-L1⁺, BCL2⁺), checkpoint⁺ (PD-L1⁺), cycling (Ki67⁺), mitotic (Ki67⁺/pHH3⁺), and immunoregulatory (IDO1⁺). Fibroblast-associated TCNs in disease were distinguished by upregulation of immunoregulatory molecules such as PD-L1, IDO1, ICOS, CD107a, and the survival factor BCL2 (Figure 7g,h).

Both survival-primed APC and immunoregulatory fibroblasts proportionally increased in disease. PD-L1 was widely expressed across fibroblast-enriched TCNs (Figure 7i). Notably, HLA-DR, HLA-A, and PD-L1 were all increased specifically in fibroblasts in periodontitis, and PD-L1⁺ fibroblasts showed strong spatial association with PD-1⁺ T cells (Figure 7j). While not an exact comparison to our transcriptomics data, some of these fibroblasts are likely some combination of stress-responsive, inflammatory, and APC-like fibroblasts defined by scRNAseq and spatial-transcriptomics data (Figure 4, 6). To test whether fibroblasts both expressed immunoregulatory ligands and engaged in spatially anchored signaling with lymphocytes, we applied receptor-ligand analysis to the spatial-proteomics dataset, focusing on the MHC-I, MHC-II and PD-L1::PD-1 checkpoint axis. This followed because an examination of our data predicted many spatially, high-confidence PD-L1::PD-1 interactions between fibroblasts (Figure 7k). As observed in the spatial-transcriptomics dataset (Figure 6), Transition zones may contain uncharacterized fibroblasts, and MCIM ligand–receptor interactions need validation beyond spatial-transcriptomics confirmation of STARComm-inferred hubs. (Figure 7l). This provides *in situ* evidence of fibroblast-driven checkpoint engagement with T cells in inflamed gingiva.

We next examined these interactions in relation to TLS-associated niches, focusing on fibroblast::T cell L::R pairs, including HLA-A⁺ fibroblasts to CD8⁺ T cells, HLA-DR⁺ fibroblasts to CD4⁺ T cells, and PD-L1⁺ fibroblasts to PD-1⁺ CD3⁺ T cells. TLS contained the highest density of these predicted interactions, with PD-L1::PD-1 signaling most frequently linking fibroblasts to regulatory T cells (Tregs), followed by CD8⁺ T cells and least to CD4⁺ T cells (Figure 7m). Given that Tregs are highly migratory and exhibit spatially dependent functional diversity, their prominence in this axis highlights the importance of considering local context when predicting fibroblast::T cell interactions. These results highlight the ability of fibroblasts to adopt multiple immunoregulatory states, positioning them as central modulators of TLS composition and function in chronic periodontitis.

## DISCUSSION (845)

This study presents the first integrated single-cell and spatial proteotranscriptomic atlas of adult human oral and craniofacial tissues, ushering in a new era of tissue-resolved mucosal immunology for the oral cavity. Building upon Human Cell Atlas principles, we developed a harmonized framework that unifies cell phenotyping, spatial communication networks, and inflammation states across oral mucosae and glands^4,55,56^. By generating standardized annotations across 13 distinct niches, we reveal how structural cells, particularly fibroblasts, spatially coordinate local immune programs. These findings expand the concept of mucosal structural immunity and highlight fibroblasts as long-lived, spatially embedded regulators of barrier-associated response.

This integrated atlas builds on our prior common coordinate framework (CCF) for the periodontium to establish a pan-oral reference map (Figure 1a)^28,57^. The CCF enables consistent identification of structural and immune cells, TCNs, and receptor-ligand networks (i.e., MCIMs), forming a scaffold for future spatial analyses and integration into the Human Cell Atlas or the Human BioMolecular Atlas Program (HuBMAP)^47,58^. As atlases expand to 2D and 3D reference structures, the ability to anchor observations to anatomical features will be critical for cross-study comparisons and biologic inference^59,60^.

A major insight from this study is the spatial anchoring of immune signaling within fibroblast-defined neighborhoods. Across mucosae and glands, fibroblasts emerged as the most transcriptionally interactive structural cell-type, exhibiting high ligand expression, spatial connectivity, and anatomical specialization. Using integrated single-cell and spatial proteotranscriptomics, we defined eight harmonized fibroblast subtypes: universal, peri-epithelial, peri-neural, peri-vascular, myofibroblasts, antigen-presenting (APC-like), immune-regulatory, and two (Type-I and -II) stress-responsive subtypes. These annotations build upon and extend fibroblast ontologies from recent integrated multi-organ atlases^21,22,24^ and by incorporating oral tissue relevant transcriptional programs and spatial localization.

Several subtypes show strong parallels with fibroblasts identified in other organs. Peri-epithelial fibroblasts, for example, resemble *COL15A1⁺* fibroblasts described in skin and gut^58^. APC-like fibroblasts share features with MHC-II⁺ populations observed in cancer^61^, synovium^62^ and cardiac tissues^63^. Fibroblasts can express MHC under inflammation, but whether this directly regulates T cells remains context-and tissue-dependent^8–10^. Stress-responsive and inflammatory fibroblasts share transcriptional signatures with those in lung fibrosis^64^, inflammatory bowel disease^65^, and synovitis^66^. Our framework refines these into a niche-aware system applicable across health and disease, supporting comparisons in conditions where fibroblast plasticity and immune regulation are altered in disease.

Fibroblast-defined MCIMs were enriched for ligands mediating cytokine *(IL1B, TNF*), chemokine (*CXCL1/2/9/10/12*), and immune checkpoint (*PD-L1, ICAM1*) signaling. These modules mapped to immune-infiltrated foci or peri-epithelial zones, echoing the concept of *para-inflammation*, a state where structural cells sustain low-level immune activity to preserve tissue homeostasis^19,20^. To spatially resolve these functional hubs, we developed a histology-2-omics (hist2omics) pipeline linking spatial-transcriptomics, multiplexed protein imaging, and same-slide histology. STARComm inferred anatomically grounded MCIMs centered on para-inflammatory or immune-active fibroblasts. More granularly, this analysis revealed fibroblast-centered therapeutic axes, including *IL1B, TNF, PD-L1*, and key chemokines, which highlight potential points of intervention across oral diseases and beyond. However, rather than pointing to a single drug readily translatable to the clinic, these findings should be viewed as a framework that nominates pathways of interest and generates hypotheses for future therapeutic exploration.

In our scRNAseq and spatial-proteomics data, fibroblasts show antigen-presenting (MHC-I/II) and immunomodulatory roles, as previously described^59–61^. In cancer, fibroblast-driven immune exclusion and checkpoint signaling suppress anti-tumor immunity, creating immunologically cold microenvironments^62–64^, and in chronic inflammation and autoimmunity^65^, modulating fibroblast-derived cues could limit tissue damage and aberrant remodeling without globally suppressing immune function^62–64^. Our atlas maps immune and fibroblast heterogeneity at single-cell and spatial resolution, enabling stromal reprogramming strategies for precision immunomodulation in cancer and chronic inflammation.^66,67^. This pan-oral proteotranscriptomic atlas lay the groundwork for perturbation-guided, fibroblast-centric therapies that reprogram stromal–immune ecosystems to durably restore barrier integrity across the body.

## Limitations of the study

Several important limitations highlight the need for continued investigation. Our dataset is primarily derived from healthy adult tissues, limiting inference in pediatric, geriatric, and disease-specific contexts. Certain niches, such as perineural spaces and ectopic lymphoid sites, were underpowered for confident annotation. Transition zones and submucosa may harbor uncharacterized fibroblast states. MCIMs and predicted ligand–receptor interactions require functional validation, as spatial-transcriptomics confirmed STARComm-inferred hubs but not receptor–ligand binding, co-localization, or downstream signaling.

Each platform has inherent limitations^68^: MERSCOPE, enables high-throughput transcript detection, but is destructive to tissue architecture, while Xenium precisely assigning transcript position relative to cell boundaries, segmentation is still a challenge. These caveats highlight the need for orthogonal approaches, including spatial-proteomics to capture protein-level signaling, histological overlays to preserve context, and *ex vivo* perturbation models (e.g., fibroblast subtype-specific targeting). Together, these complementary strategies will provide a more comprehensive and functionally grounded view of multicellular communication..

In summary, we present a comprehensive single-cell and spatial proteotranscriptomic atlas of adult human oral tissues, a harmonized fibroblast classification system, even presenting two stress-responsive fibroblast states hereunto undescribed in literature and influenced by their niche-of-origin, as well as modular tools to interrogate spatially resolved immunoregulation. By integrating histology, spatial multiomics, and computational inference, we lay the groundwork for targeted modulation of fibroblast activity in fibrosis, cancer, and autoimmunity. This foundational map enables future efforts to model, compare, and therapeutically manipulate structural–immune circuits across development, aging, and disease.

## SUPPLEMENTARY DATA LEGENDS

**Supplementary Figure 1:**
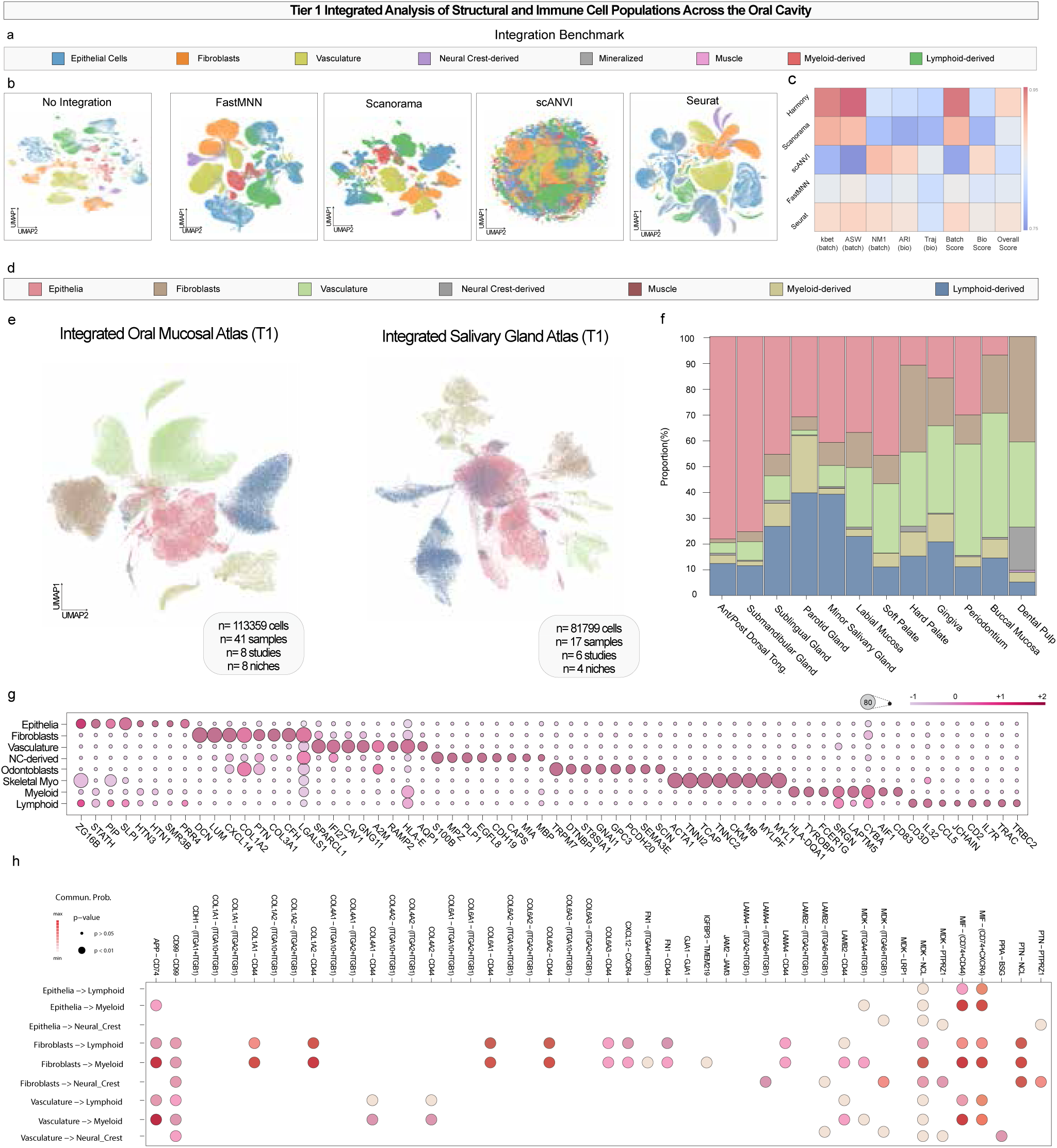
Niche Diversity and Cell Types within the Oral and Craniofacial Atlas. (a-c) Integration of OCF data using (a) Tier 1 ontologies and (b) benchmarking single-cell integration using (c) five batch-correction methods (Harmony, Scanorama, BBKNN, FastMNN, scVI) evaluated with scIB metrics. (d,e) Integrated scRNA-seq Tier 1 atlas of oral mucosae and salivary glands using (d) Tier 1 ontologies subclustered into (e) glands and oral mucosae (all available via CELLxGENE). (f) Cell type proportions within each integrated dataset. (g) Differentially expressed genes (DEGs) for Tier 1 cell type annotations across integrated datasets. (h) CellPhoneDB analysis showing significant ligand–receptor interactions between structural and immune cell types.

**Supplementary Figure 2:**
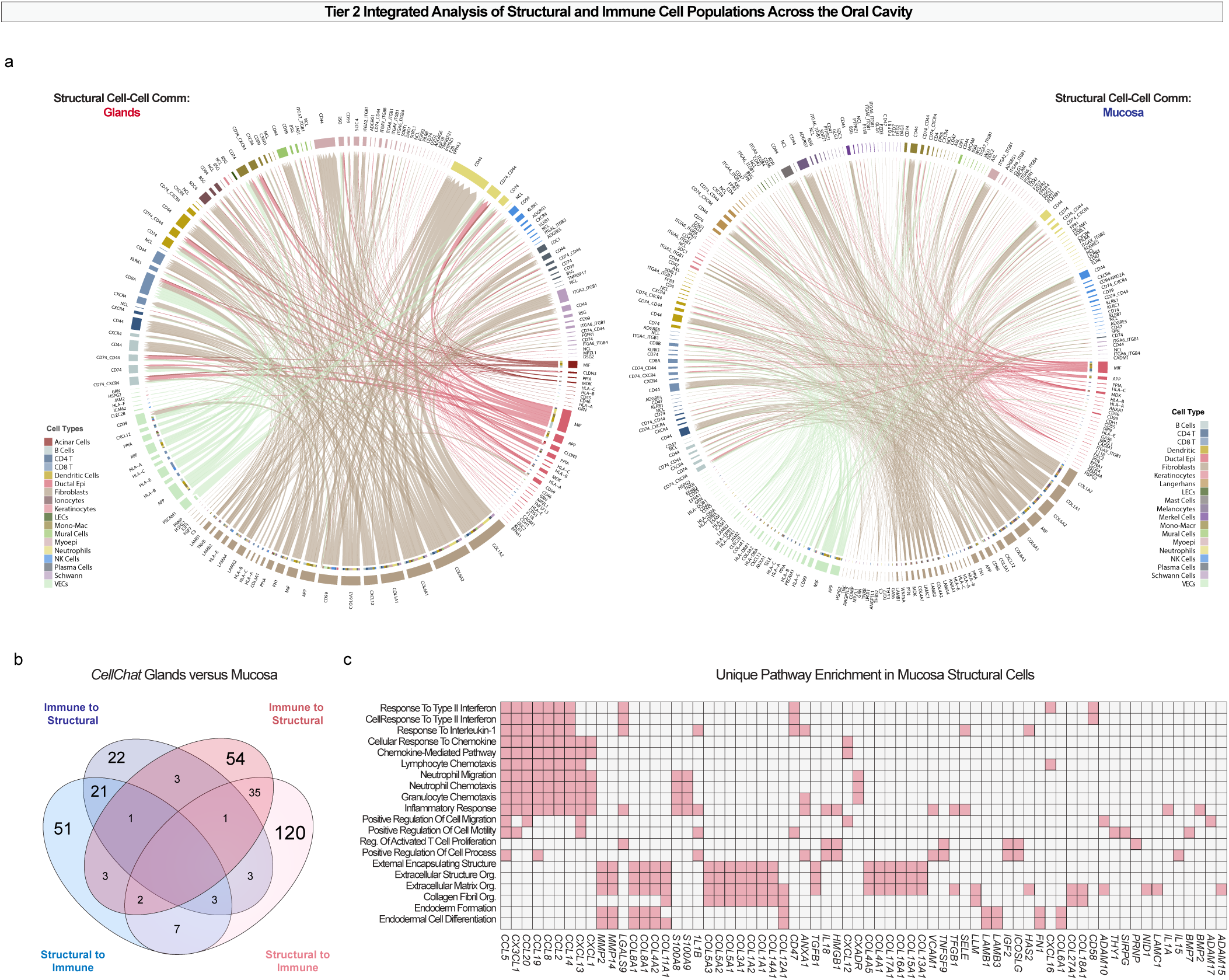
Cell::cell communication patterns of structural cell types in glands and mucosae. (a) Chord diagrams depicting structural cell–cell communication in glands (left) and mucosae (right), highlighting inferred ligand–receptor interactions between structural and immune cell populations. (b) Venn diagram comparing the overlap of ligand– receptor pairs between glands and mucosae, stratified by directionality of communication: immune-to-structural and structural-to-immune. (c) Heatmap showing unique pathway enrichment in mucosal structural cells, with emphasis on pathways involved in innate immune cell recruitment. These pathways contribute substantially to predicted structural::immune interactions, underscoring their role in mucosal tissue homeostasis and in the initiation and progression of disease-associated processes.

**Supplementary Figure 3.**
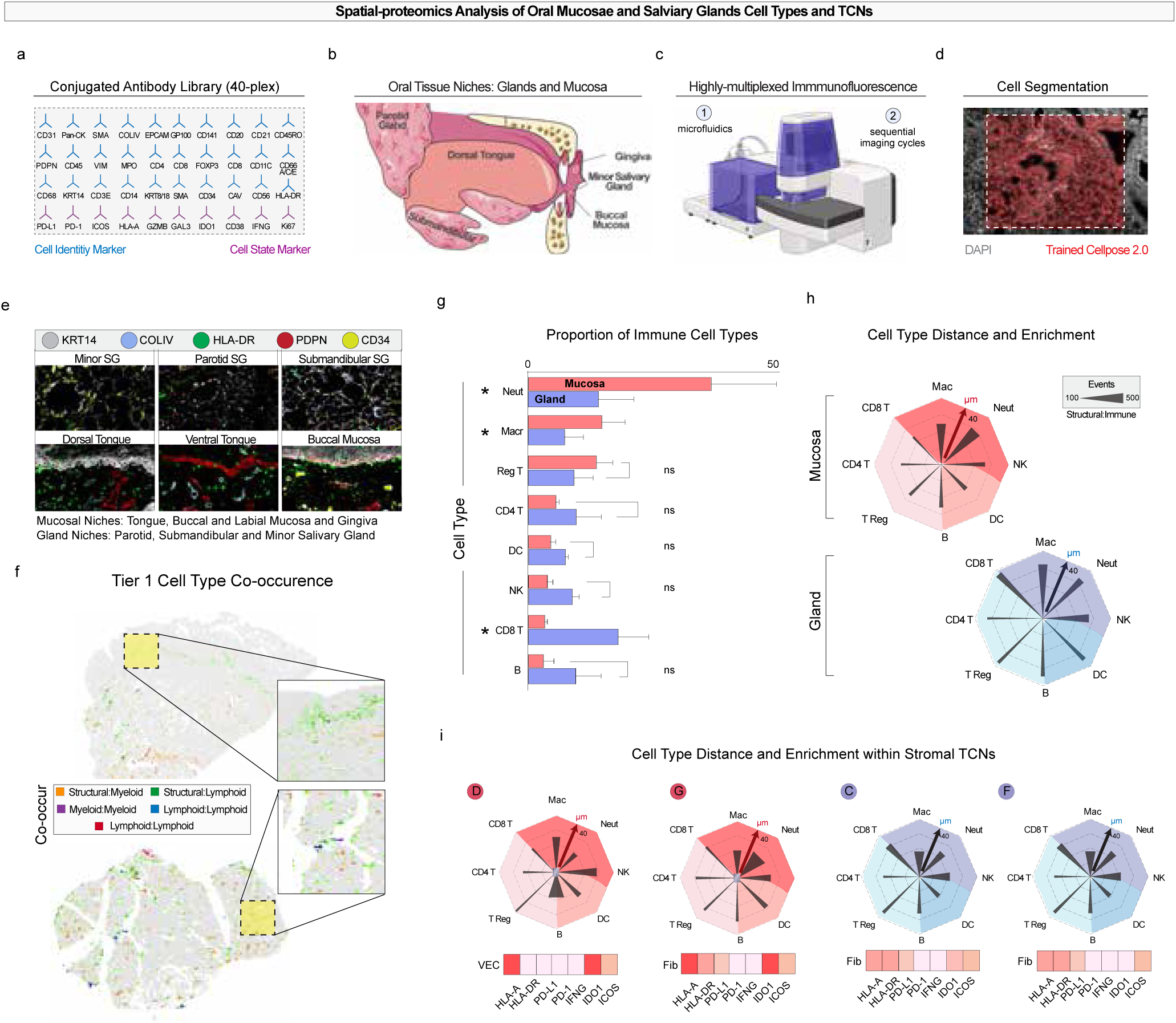
Building and validating a spatial-proteomics atlas of the oral cavity. (a–c) Using Phenocycler Fusion technology, we developed a 40-plex oligo-conjugated antibody panel (a) to analyze the tropism and colocalization of innate and adaptive immune cells within specific neighborhoods and structures across six distinct oral tissues (b), processed two slides at a time (c). (d) To extract single-cell spatial data, we trained a segmentation model on oral cavity tissues using Cellpose 2.0. (e) Representative multiplex immunofluorescence images from mucosa and salivary glands, showing selected markers from the PCF panel. (f) Tier 1 co-occurrence maps highlight structural–myeloid, structural–lymphoid, myeloid–myeloid, lymphoid– myeloid, and lymphoid–lymphoid interactions across tissues. (g) Quantification of immune cell proportions in mucosal versus glandular niches (ns, not significant). (h) Pairwise distance and enrichment analyses reveal preferential proximities between structural and immune subsets in mucosal (top) and glandular (bottom) tissues. (i) Heatmaps showing cell type enrichment, immunophenotypes, within stromal tissue cellular neighborhoods (TCNs). Statistical test: one-way ANOVA and t-test, FDR-adjusted p < 0.05.

**Supplementary Data 4.**
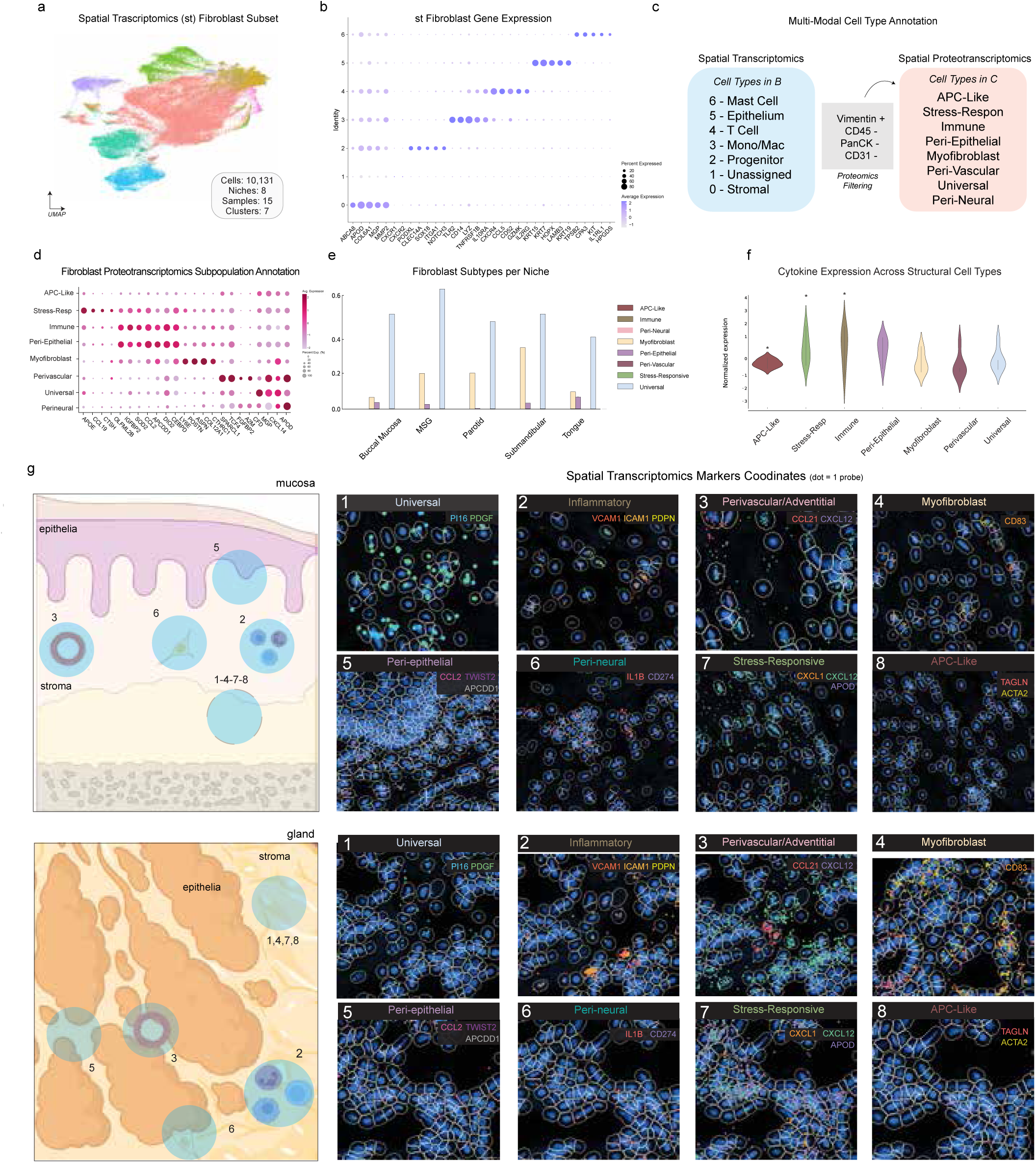
Fibroblast subpopulation annotation and spatial localization in oral mucosae and salivary glands. (a) UMAP of all fibroblasts after protein-anchored filtering for VIM+ cells. (b,c) Clusters of Fibroblasts as predicted by spatial-transcriptomics. Which when annotated show mixed cell types (c) that are clarified into the scRNAseq cell types predicted in Figure 4 only after using protein anchoring. (d) Dot plot showing fibroblast subpopulation annotation across tissue niches, defined by single-cell RNA-seq and spatial-transcriptomics integration. Size of the dot indicates the proportion of cells, and color intensity corresponds to scaled average expression of subpopulation marker genes. (e) Relative abundance of fibroblast subtypes across structural niches, quantified as a fraction of total fibroblasts within each tissue region. (f) Violin plots of cytokine expression across structural cell types, showing niche-specific transcriptional programs. (g) Schematic illustrations (left) depict epithelial and stromal compartments of oral mucosa (top) and salivary gland (bottom), with colored circles corresponding to fibroblast subpopulations identified in (a–c). Spatial-transcriptomics maps (right) display representative fields for each fibroblast subpopulation: Universal (1), Inflammatory (2), Perivascular/Adventitial (3), Myofibroblast (4), Peri-epithelial (5), Peri-neural (6), Stress-Responsive (7), and APC-Like (8). Colored outlines indicate segmentation of cell boundaries; marker genes used for subpopulation identification are labeled in each panel. Abbreviations: APC, antigen-presenting cell; GO, Gene Ontology; FDR, false discovery rate. Scale bar (g):100 µm

**Supplementary Figure 5.**
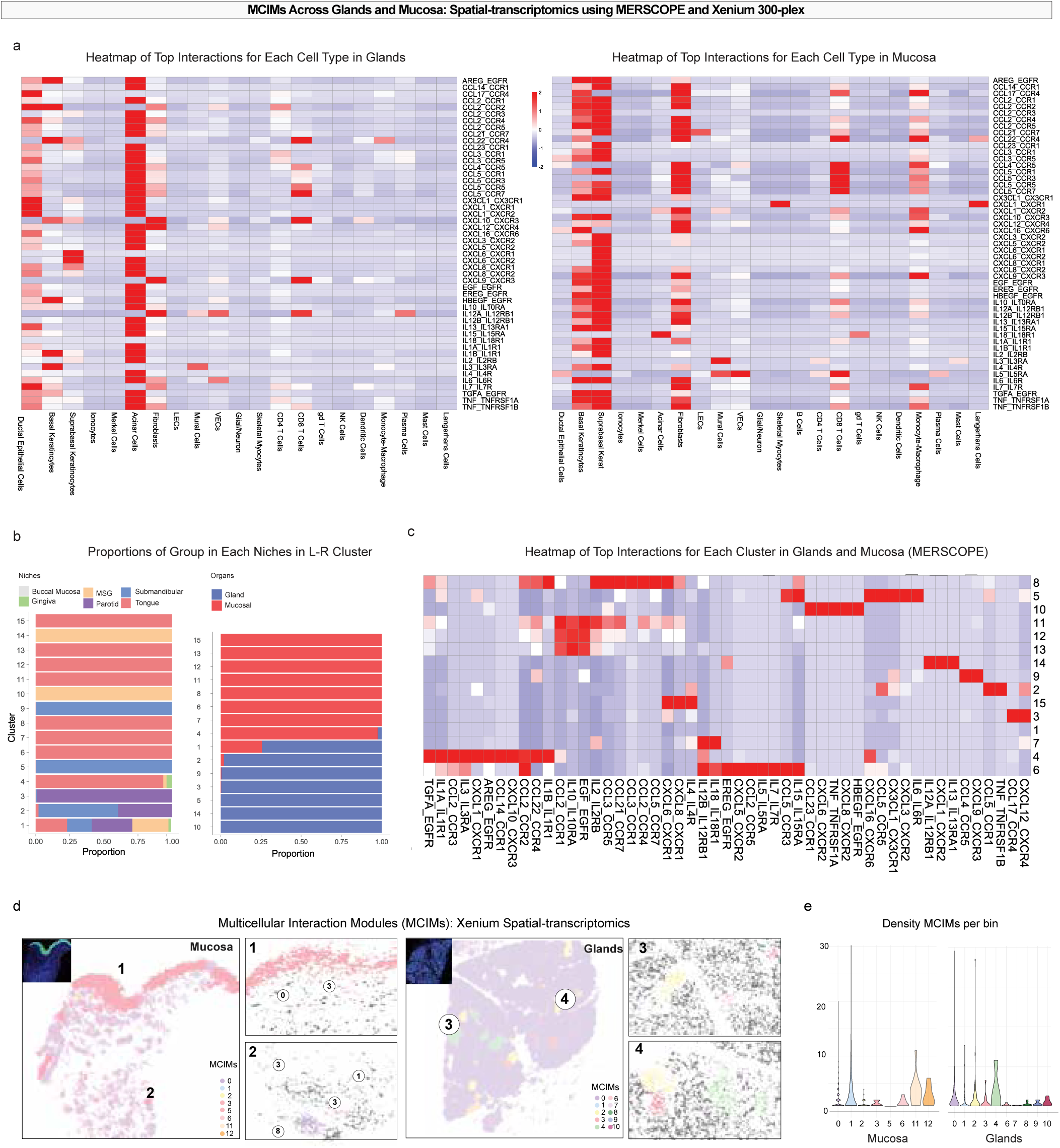
MCIMs Across glands and mucosa using spatial-transcriptomics. (a) Heatmaps of the top ligand–receptor (L–R) interactions for each cell type identified in glands (left) and mucosa (right) using MERSCOPE spatial-transcriptomics. Interaction scores are shown with red indicating higher and blue indicating lower predicted interaction strength. (b) Proportions of regional/niche-specific clusters from L–R analysis, grouped by organ type (gland vs mucosal) and specific niche (buccal mucosa, gingiva, minor salivary gland [MSG], parotid, submandibular, tongue). (c) Heatmap of the top L–R interactions for each R–L cluster in glands and mucosa (MERSCOPE). (d) Representative examples of multicellular interaction modules (MCIMs) identified in mucosa (1–2) and glands (3–4) using Xenium 300-plex spatial-transcriptomics. (e) Plots showing MCIM density per spatial bin for mucosa (left) and glands (right). Abbreviations: L– R, ligand–receptor; MCIM, multicellular interaction module; MSG, minor salivary gland.

**Supplementary Figure 6.**
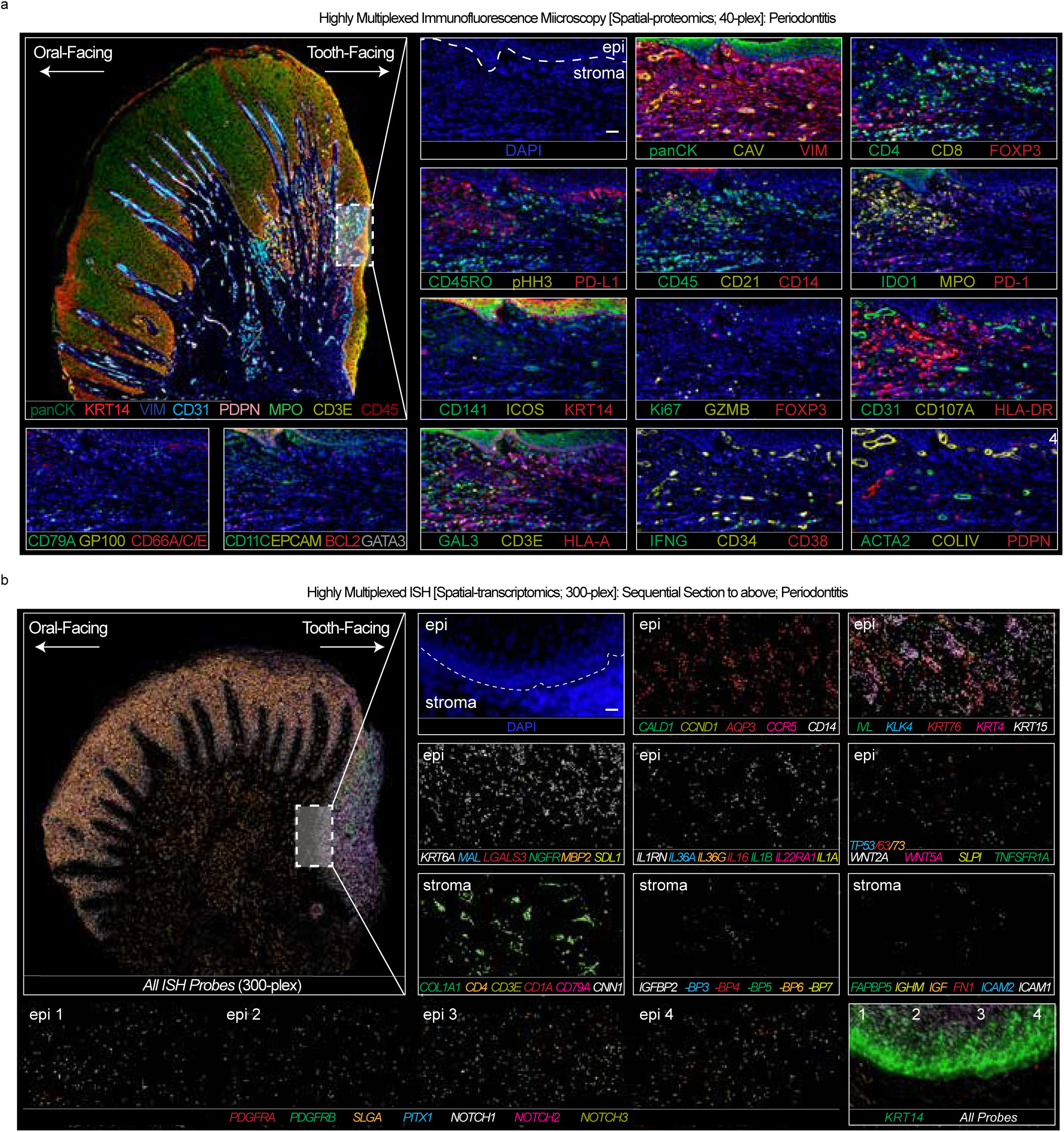
Spatial Proteomic and Transcriptomic Mapping of Gingiva in Periodontitis. | Representative spatial-proteomics and spatial-transcriptomics images of oral mucosal tissues, showing region-specific protein and gene expression patterns. (a) Spatial-proteomics images acquired using the Akoya Phenocycler-Fusion platform, stained with the OMAP-29 antibody panel. (b) Spatial-transcriptomics images generated with the 10x Genomics Xenium platform, showing cell-type-specific transcript localization. Insets highlight areas of interest for immune–stromal–epithelial interactions. Scale bars = 100 µm for all panels.

**Supplementary Figure 7.**
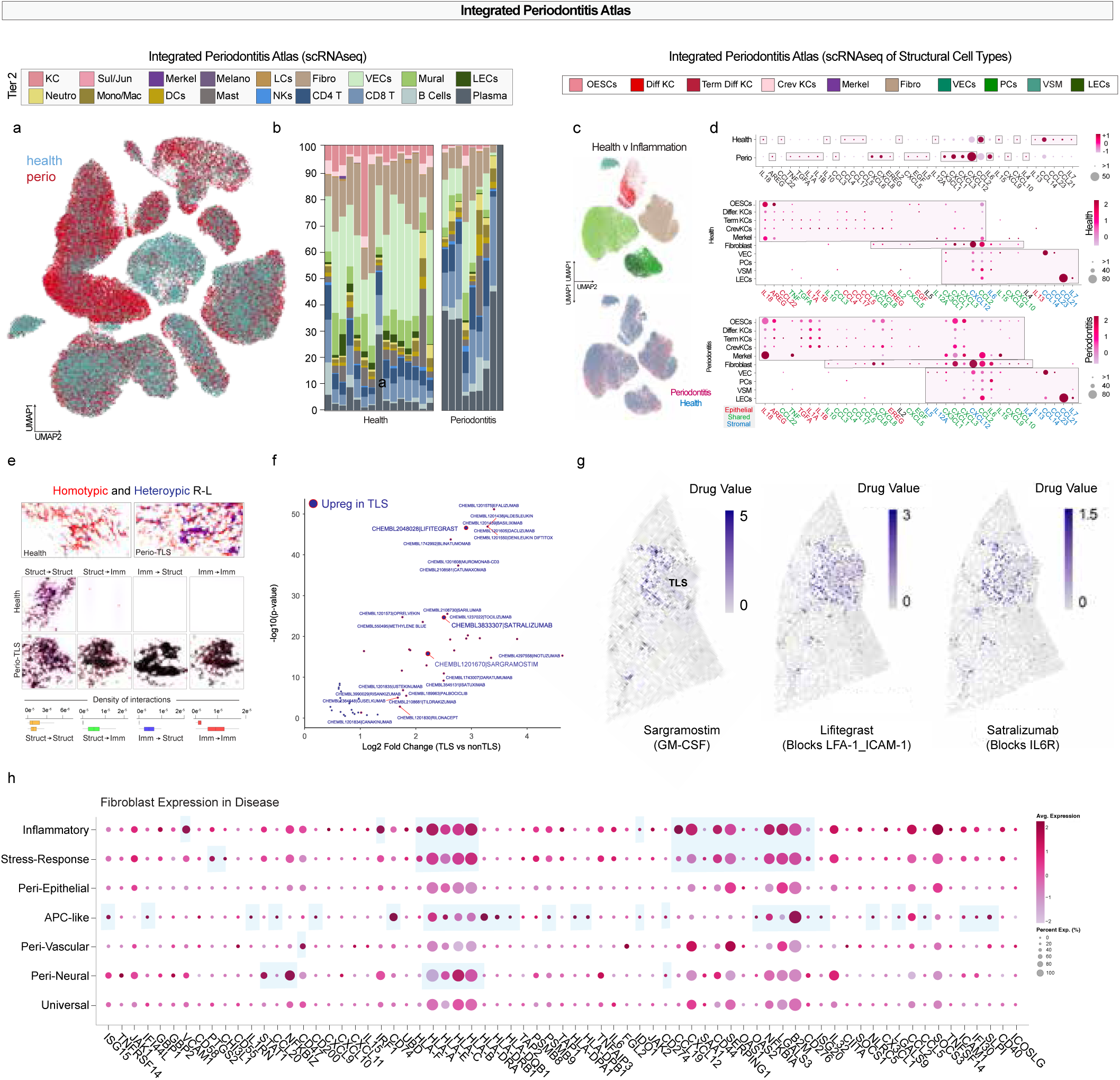
Integrated Periodontitis Atlas across health and disease states with ligand–receptor interaction and drug targeting analysis. (a) UMAP projection of integrated single-cell RNA sequencing (scRNA-seq) datasets from healthy (blue) and periodontitis (red) gingival tissues, annotated by immune and stromal cell identities (Tier 2 annotation). (b) Stacked bar plots showing proportional representation of immune and stromal cell types in healthy and periodontitis samples. (c) UMAP projection of integrated scRNA-seq datasets subset to structural cell types, colored by disease status (health vs periodontitis). (d) Dot plots showing differentially expressed ligands and receptors between healthy and periodontitis within structural cell types, with dot size representing percentage of cells expressing the gene and color indicating average expression. (e) Representative spatial transcriptomic (spRNA-seq) maps and density plots illustrating homotypic and heterotypic ligand–receptor (R–L) interactions between structural and immune cells in health and in tertiary lymphoid structures (TLS) during periodontitis. (f) Volcano plot showing ligands and receptors upregulated in TLS compared to non-TLS regions, highlighting chemokine signaling molecules. (g) Spatial drug value maps illustrating predicted targets of TLS-upregulated ligand–receptor interactions for candidate therapeutics with color intensity corresponding to relative drug value scores. (h) Fibroblast expression only in disease across the newly annotated subtypes shows evidence for linked immunoregulatory signaling within and across cell states.

## METHODS

### Published scRNAseq harmonization

Raw FASTQ files from published single-cell RNA sequencing projects were retrieved using scripts from [this GitHub repository](https://github.com/cellgeni/reprocess_public_10x). First, metadata was collected from the GEO soft family file, and the ENA web API was used to gather data format details (SRR/ERR) and link samples to runs. Raw reads were downloaded as SRA archives, 10X BAM files, or gzipped paired-end FASTQ files, and then converted to FASTQ using fastq-dump (SRA tools v2.11.0) or bamtofastq (v1.3.2). The raw reads were processed using STARsolo for mapping and quantification, with wrapper scripts from https://github.com/cellgeni/STARsolo/ to identify kit versions and sample specifics. The human reference genome matched Cell Ranger 2022-A standards. For 10x samples, STARsolo settings were optimized to mimic Cell Ranger v6 outputs, including UMI/barcode processing and paired-end alignment. Cell filtering was performed with EmptyDrops (Cell Ranger v4+), producing both exon-only and full-length gene counts as well as RNA velocity matrices.

### Newly generated scRNAseq data

*<u>Soft Palate (MSNZ Würzburg)</u>*: Resected soft palate oral mucosa samples were placed in ice-cold Advanced DMEM+++ medium (Advanced DMEM F12 (Gibco) supplemented with 10 mmol/l HEPES (Gibco), 100 U/ml penicillin/streptomycin (Gibco), 1 x GlutaMax (Gibco), and 10 µM of the Rho kinase (ROCK) inhibitor compound Y-2763) and transferred to the laboratory on ice as soon as possible. For histology, representative pieces of tissue were embedded in Tissue-Tek® O.C.T. compound (Sakura) and placed directly on dry ice. The O.C.T.-embedded tissues were stored at –80°C for long-term storage. O.C.T. blocks were cut into 10 mm sections and stained with hematoxylin and eosin (H&E, Morphisto) after PFA fixation. For cryopreservation, the tissues were cut into smaller pieces and were incubated in CryoStor® CS10 (Stem Cell Technologies) freezing medium for 10 minutes on ice. Then, the samples were transferred to a –80°C freezer. Frozen samples were stored in –150°C long-term.

Frozen tissue samples were thawed at 37°C and washed in pre-warmed Advanced DMEM+++ medium. After rinsing, the tissue pieces were incubated in Advanced DMEM+++ for 10 min at room temperature. Samples were minced to a size of 0.5–1 mm3 and digested in Advanced DMEM+++ supplemented with 2% fetal bovine serum (Gibco), 2 mg/ml collagenase-P (Sigma-Aldrich), 1 mg/ml collagenase-D (Sigma-Aldrich), 0.5 mg/ml DNase-I (Sigma-Aldrich) and 10 µmol/ml Y-27632 (Tocris) for 30 minutes in a thermomixer (Eppendorf) at 37 °C, 1000 rpm. During incubation, the digestion mixture was vigorously resuspended every 10 minutes. Samples were further digested into single cells in 0.2% trypsin in PBS supplemented with 10 µmol/mL Y-27632 (Tocris). The digestion was stopped with EDTA. Single cell suspension was filtered through a pre-wetted Pluristrainer with a pore size of 70 µm. Antibody staining and sorting: Cells were stained with Zombie Violet Fixable Viability Stain (1:1000 in PBS, Thermo Fisher) and blocked with Human TruStain FcX™ Fc Receptor Blocking Solution (1:10 in PBS, Biolegend). Then, cells were stained with TotalSeqTM-B0251 and B0252 (1:50, Biolegend), and alive cells were sorted with a BD FACS Aria III flow cytometer and collected into a clean tube. Sorted samples were pooled, counted with Trypan Blue, and loaded onto a 10x Chromium Controller (10x Genomics). Gene expression and cell surface protein library preparation was performed according to the manufacturer’s instructions for the 10x Chromium Next GEM Single Cell Library kit v3.1 Dual Index (10x Genomics). The libraries were sequenced on a NextSeq 2000 Illumina sequencer. *<u>Labial Mucosa (National Institutes of Health):</u>* Samples were collected from patients who provided informed consent under the NIH Central IRB Protocol 15-D-0051 (Principal Investigator: Warner). The sections were immediately placed in ice-cold RPMI, dissected into 1–2-mm pieces, and dissociated using a Miltenyi Multi-tissue Dissociation Kit A. It was then placed in 10% formaldehyde for a 16-24h fixation at 4°C during the fixation step and subsequently placed in a - 80 fridge. To initiate the thawing process, the Thaw Enhancer (10x Genomics PN-2000482) was incubated at 65°C for 10 minutes, followed by vortexing and a brief centrifugation to ensure no precipitate was present. The reagent was kept warm, and the absence of precipitate was verified before use. Thawed Enhancer was not kept on ice to prevent precipitation, and once thawed, it was maintained at 42°C for up to 10 minutes. Next, 0.1 volume of pre-warmed Enhancer was added to the fixed sample in Quenching Buffer. For example, 100μl of Enhancer was added to 1,000μl of fixed sample in Quenching Buffer, followed by pipetting to mix. Alternatively, to conserve Enhancer volume, cells were centrifuged at 850 RCF, 500μl of Quenching Buffer was removed, and 50μl of Enhancer was added to the sample. 50% glycerol was introduced to achieve a final concentration of 10%. For instance, 275μl of 50% glycerol was added to 1,100μl of fixed sample in Quenching Buffer and Enhancer, followed by pipetting to mix. The samples could be stored at -80° for up to 6 months. In the post-storage processing phase of our study, all the samples were processed at the same time and, the steps outlined below were followed: Samples were thawed at room temperature until no ice was present. The samples were then centrifuged at 850rcf for 5 minutes at room temperature. The supernatant was carefully removed without disturbing the pellet. The cell pellet was resuspended in 1 ml of 0.5X PBS + 0.02% BSA* (optionally supplemented with 0.2 U/μl RNase Inhibitor) or Quenching Buffer and kept on ice.

*RNase-free BSA was used at this step. The dissociation process was conducted at 37 °C in an OctoMACS tissue disruptor using heated sleeves. Single-cell suspensions underwent serial filtration through 70- and 30-µm filters and were rinsed with 1× Hanks’ buffered salt solution. Cell counting and viability assessments were performed using a Trypan blue exclusion assay, with suspensions having greater than 35% viability chosen for subsequent sequencing. For single-cell capture, library preparation, and sequencing, approximately 10,000 cells were targeted and loaded onto a 10x Genomics Chromium Next GEM Chip B. Following cell capture, library preparation was performed using the 10x Chromium Next GEM Single Cell 3′ kit v3, and the libraries were sequenced on a NextSeq500 sequencer (Illumina). Cell processing was undertaken using the 10x Genomics Chromium Controller and the Chromium Single Cell 3′ GEM, Library & Gel Bead Kit v3 (PN-1000075), adhering to the guidelines provided in the manufacturer’s user guide. Viability and concentration assessments were conducted by staining aliquots of sorted cells with acridine orange and propidium iodide, followed by analysis using a fluorescent Cell Counter. Each sample was loaded with approximately 3,200 cells onto the Chromium Chip B, aiming for a recovery of 2,000 cells per sample during library preparation. The creation of gel beads in emulsion (GEMs) involved encapsulating single cells, reverse transcription reagents, and gel beads coated with barcoded oligos within an oil droplet. Reverse transcription was executed using a C1000 thermal cycler (Bio-Rad), resulting in complementary DNA (cDNA) libraries tagged with a cell barcode and unique molecular index (UMI). GEMs were then broken, and the cDNA libraries purified using Dynabeads MyOne SILANE (Invitrogen) with subsequent 12 amplification cycles. Amplified libraries underwent purification with SPRIselect magnetic beads (Beckman Coulter) and quantification using an Agilent Bioanalyzer High Sensitivity DNA chip (Agilent Technologies). Following steps included fragmentation, end repair, A-tailing, and double-sided size selection with SPRIselect beads. Ligation of Illumina-compatible adapters onto size-selected cDNA fragments occurred. Adapter-ligated cDNA was purified using SPRIselect beads, and uniquely identifiable indexes were incorporated during 12 amplification cycles. The finalized sequencing libraries underwent purification with SPRIselect beads, visualization using the Bioanalyzer High Sensitivity DNA chip, quantification with the KAPA SYBR FAST Universal qPCR Kit for Illumina (Roche) and StepOnePlus Real-Time PCR System (Applied Biosystems), normalization to 4nM, and pooling. Sequencing took place on a NextSeq 500 machine (Illumina), with libraries denatured and diluted following the standard Illumina protocol. A 1% PhiX sequencing control (Illumina) was spiked in, and the libraries were loaded onto the flow cell at 1.8pM. *<u>Parotid Salivary Glands (University of Groningen):</u>* Biopsy samples of parotid SG tissue were obtained from donors after institutional review board [IRB] approval, (METc 2008.078, METc 2023.281) who were treated for a squamous cell carcinoma of the oral cavity. In these patients, an elective head and neck dissection procedure was performed. During this procedure, the parotid SG is exposed and removed. This tissue does not contain malignant cells, as oral squamous cell carcinoma does not disseminate to the parotid SG. Salivary gland tissue was processed into a cell suspension according to our previously published method (PMID: 29984480), and CS10 cryopreservation media until use. After thawing, separate cells were processed using 10X Genomics protocol version 3.1 chemistry (PN 2000121) accompanying the Version 3.1 kit, with a targeted cell yield of 8000 per lane. Samples were not multiplexed per 10X machine lane. Sample indexes from the Chromium i7 Sample Index Kit (PN-120262) were used for identification of pooled sampled on sequencing chip. Sequencing was performed using the Next Seq2500 (RapidRun) sequencer for a total of 127 cycles.

### Published scRNAseq Integration, pre-process and visualization, and spatial single-cell annotation using Trailmaker

*<u>scRNAseq:</u>* The single-cell RNA sequencing dataset was analyzed and visualized using Trailmaker®, which was first accessible on the Biomage community platform (Biomage; https://biomage.net/), now developed and maintained by Parse Biosciences. Pre-filtered count matrices were loaded onto the platform for additional processing. Barcodes were filtered in four steps: first, barcodes with fewer than 500 UMIs and those indicating low quality or signs of apoptosis (over 15% mitochondrial content) were excluded. A robust linear regression model from the ‘MASS’ package was then used to predict expected gene counts per barcode, applying a tolerance based on sample size to exclude outliers outside this range. Doublets were identified and removed using the ‘scDblFinder’ R package, filtering out barcodes with a score over 0.5. Following filtering, between 300 and 8,000 barcodes per sample remained. Data was normalized with a log transformation, and the top 2,000 most variable genes were identified using variance-stabilizing transformation. PCA was applied, with the first 40 components (capturing 95.65% of the variance) then batch-corrected using the ‘Harmony’ package. To benchmark data integration, we applied the scIB pipeline to evaluate five methods: Harmony, Scanorama, scVI, BBKNN, and FastMNN. Integration performance was assessed across both batch correction and biological conservation. For batch correction, we evaluated overall performance using principal component regression (PCR), average silhouette width (ASW), and graph-based iLISI scores. For biological conservation, we used cell cycle variance and normalized mutual information (NMI) to assess the preservation of biological structure. Louvain clustering and UMAP were conducted in Seurat for dimensionality reduction and cluster identification. Differentially expressed genes were identified for each cluster using the Wilcoxon rank-sum test (‘prestò package). These samples were then imported into Trailmaker®, bypassing initial filtering since pre-processing had been completed. The same analytical pipeline was applied to this subset as to the complete dataset. Cells were annotated based on literature and further classified with the CellTypist tool. For MERFISH data from MERSCOPE, segmented data, including barcodes, features, and matrices, was processed within Trailmaker®. Cells with no transcripts were removed based on transcript count per cell, and cell types were annotated using the lasso tool, focusing on the most highly expressed genes within each Louvain cluster.

### Receptor-ligand analyses via *Cellphone DB*^67^, *CellChat*^68^, and *MultiNicheNe*t^69^

We employed *CellPhoneDB* (v5, https://github.com/ventolab/CellphoneDB), leveraging its curated database of ligand-receptor interactions to analyze cell-to-cell signaling interactions at a single-cell level. Using default parameters, we calculated the likelihood of interactions between pairs of cell types based on the expression of known ligand-receptor pairs, resulting in probabilistic scores that highlight statistically significant intercellular communication. Additionally, we applied CellChat (v2) to quantify signaling probabilities between manually defined sender and receiver cell groups. Both methods were applied to the same datasets, providing a comprehensive communication profile across different niches and enabling cross-validation of key interactions. We also employed *MultiNicheNet*, a novel framework available at https://github.com/saeyslab/multinichenetr, to enhance the analysis of cell-cell communication within multi-sample multi-condition single-cell transcriptomics data, with a focus on oral mucosal tissues. The primary objectives of MultiNicheNet were to infer differentially expressed and active ligand-receptor pairs between conditions of interest and predict the putative downstream target genes of these pairs. The pipeline was applied in two different approaches, one related to a Tier 1 annotation dataset into the integrated oral craniofacial atlas. Statistical difference was employed to determine the top 50 interactions between receptor and ligands in glands versus mucosa. A second approach was made to determine the difference within each of the major niches, such as Salivary Glands and all Mucosa. Cells annotations were carried from Tier 2 to run downstream analysis. In the purpose of setting the threshold of cell, only cell types with more than 10 cells in all niches were included in the analysis. p-values were extracted using muscat framework for DEG and testing which is the top highest expressed based on mixed linear models.

### Sectioning for spatial biology

Samples from the pathology archive of UNC medicine school (UNC IRB 22-1786) were selected with two trained oral pathologists, only samples not containing architectural or cytologic alterations were included. FFPE Blocks were sectioned in 5 microns sequential sections and mounted in SuperFrost Plus (Thermo Fisher) slides for Phenocycler-Fusion 2.0, H&E and Stains. Coverslips (MERSCOPE slides) from MERSCOPE (Vizgen) were also used to mounting the sequential sections from each slide that went through other Phenocycler experiment using the ‘91600112 MERSCOPE User Guide Formalin-Fixed Paraffin-Embedded Tissue Sample Preparation RevB’. MERSCOPE TM Slides have a defined imageable of a 2X1.5cm rectangular where two sections from the same tissue were placed at time. All slides preparation was performed using RNA free water and RNAse protocol from Leica autotec system.

### Multiplex Protein Immunofluorescence (Formerly, CODEX; now, PhenoCycler Fusion 2.0, Akoya Biosciences)

For multiplexed immunofluorescence (Multi-IF), we used the PhenoCycler Fusion 2.0 from Akoya Biosciences. All samples underwent deparaffinization using a gradual reduction of alcohol concentrations from 100% to 30%. Antigen retrieval was performed using AR9 (EDTA) buffer from Akoya Biosciences in a low-pressure cooker for 15 minutes. After cooling for 1 hour, samples were hydrated with ethanol for 2 minutes followed by a 20-minute incubation in staining buffer. The antibody cocktail buffer was prepared according to the Phenocycler Fusion Manual using four blockers and nuclease-free water in addition to the staining buffer. Antibodies (Supplementary Table 6) were diluted to a 1:200 concentration in the antibody cocktail buffer. A mixed solution containing all primary antibodies was then prepared. Slides were incubated overnight at 4°C in a humidity chamber (Sigma-Aldrich) with the primary antibodies. After primary incubation the slides were placed in staining buffer for 2 min, followed by post-stain fixing solution (10% PFA in staining buffer) for 10 min. Following 3 baths of 2 min washing in 1X PBS, the slide was immersed in ice-cold MeOH for 5 min. The final fixative solution (FFS) was prepared in accordance with the Phenocycler manual. Slides were then incubated with the FFS for 20 minutes at room temperature. Following this, the slides underwent a PBS wash and were promptly transferred to the FCAD Machine (Akoya Biosciences) for flow cell mounting. Flow cell stitching was achieved through a 30-second application of high pressure to the top of the slides. Once mounted, the flow cells with the slides and sectioning were immersed in PhenoCycler (PCF) buffer for 10 minutes before being transferred to the Phenocycler Fusion. Reporters for each antibody were prepared using a Report Stock solution along with 5490 µL of nuclease-free water, 675 µL of 10X PCF buffer, 450 µL of PCF assay reagent, and 27 µL of in-house prepared concentrated DAPI to achieve a final DAPI concentration of 1:1000 per cycle. This process was repeated for a total of two slides per time. Reporters were included into a 1:50 dilution for each cycle, including the respective channels. 250 microliters of each specific report stock solution were pipetted into a 96 well-plate that was sealed using an aluminum foil provided by Akoya Biosciences. Two slides were included into the fluid equipment at time, using the Phenocycler Fusion 2.0. Manual mapping was used to the scanning area to be scanned by the PhenoImager using brightfield. Low DMSO and High DMSO was prepared as suggested by the Index B from the Phenocycler Fusion 2.0 manual of operations.

### Multiplexed error-robust fluorescence in *situ* hybridization (Vizgen MERSCOPE)

For multiplexed error-robust fluorescence *in situ* hybridization (MERFISH), we used MERSCOPE from Vizgen. To ensure maximum sensitivity and consistency, MERSCOPE verification was performed on representative samples from all oral niches using the MERSCOPE Verification Kit (Vizgen) according to the manufacturer’s instructions (91600004 Rev D). Briefly, verification mimics all sample processing steps but uses a single-round smFISH readout of the EEF2 housekeeping gene to determine signal brightness, background fluorescence and consistent adhesion. Verification results with increased background in several tissue types made alterations to the manufacturer’s protocol necessary. A harsh clearing protocol was developed and verified on additional samples in alignment with the Vizgen team. It replaced the usual digestion followed by single stage clearing of clearing resistant tissues with a multi-step protocol with increased Proteinase K (ProtK, NEB, Cat P8107S) concentrations. After gel embedding, samples were initially cleared with 1.25 ml clearing premix (Vizgen) and 120 µl ProtK for 4 hours at 47 °C. After incubation another 3.75 ml of clearing premix without ProtK was added to the samples and incubated over night at 47 °C. Samples were washed three times for 5 minutes in Sample Prep Wash Buffer (Vizgen) to remove the clearing solution and digested in Digestion Mix (Vizgen) for 2 hours at 37 °C. 5 ml fresh Clearing Premix with 50 µl ProtK were added directly to the digestion reaction and incubated for 3 days at 37 °C before continuing with autofluorescence quenching according to the unmodified MERSCOPE protocol. These modifications to the FFPE MERSCOPE protocol (91600112 Rev B) were used to generate all MERSCOPE results otherwise following the manufacturer’s instructions. Samples were hybridized with MERSCOPE probes for the RNAs listed in Supplementary table 5 and smFISH probes for KRT14 (ENSG00000186847), KRT1 (ENSG00000167768), KRT6B (ENSG00000185479), APOD (ENSG00000189058), LORICRIN (ENSG00000203782) and B2M (ENSG00000166710) that were all part of the custom designed panel (Vizgen panel ID: BP0983). Next to transcript locations, tiff images were generated for DAPI and polyT as well as three proprietary cell boundary stains (Vizgen, PN 10400009) at 0.108 µm pixel size and the smFISH probes listed above.

### Xenium Spatial-transcriptomics and Histo2mics

For spatial transcriptomic profiling, FFPE tissue sections were processed using the Xenium *in Situ* platform (10x Genomics) according to the manufacturer’s user manual. A custom-designed probe panel targeting 300 genes of interest was used for transcript detection. Tissue sections were mounted onto Xenium-compatible slides and underwent the full automated workflow, including tissue permeabilization, probe hybridization, and signal amplification, as outlined in the protocol. Following the completion of the Xenium run, we opted not to proceed with the standard hematoxylin and eosin (H&E) staining. Instead, we performed the optional dequenching step described in the Xenium protocol, which enables removal of residual fluorescent signal and preserves the tissue for subsequent downstream applications, such as multiplexed spatial-proteomics. Following the dequenching step, the slides were placed in a storage solution consisting of 10% glycerol and 90% phosphate-buffered saline (PBS) to preserve tissue integrity and morphology. Slides were stored at 4°C in this buffer until further processing for spatial-proteomics, ensuring compatibility with downstream multiplexed immunofluorescence workflows. Slides were then subjected to a new round of antigen retrieval using EDTA-based buffer pH 9 (Akoya Biosciences) to prepare the tissue for multiplexed immunofluorescence. The same Phenocycler panel used in subsequent experiments was applied to ensure consistency across samples. After completion of the spatial-proteomics workflow, slides were immersed in HistoChoice® MB Tissue Fixative (VWR International, Radnor, PA) for 24 hours to facilitate flow cell removal. The flow cell was carefully detached using a #15 surgical blade, taking care not to damage the underlying tissue. Finally, slides were stained with hematoxylin and eosin (H&E) to assist in tissue segmentation and provide morphological context for downstream analyses. The final output consisted of three spatially aligned images obtained from the same slide: Phenocycler multiplexed proteomics, Xenium spatial-transcriptomics, and H&E histology.

### Cell Segmentation for Multi-IF and Spatial-transcriptomics Images using *Cellpose3*^48^

Cell segmentation was carried out using *Cellpose3*. We employed a pipeline that integrated both denoising and segmentation steps to improve the quality of input images and the accuracy of the segmentation outputs based in nuclei expansion. The model was custom-trained on H&E-stained tissue sections from the niches of the oral cavity, allowing for optimized performance on this tissue type, which often presents significant histological heterogeneity. Prior to segmentation, a denoising step was applied to reduce image noise and artifacts. The cell expansion was based on the auto-calibration tool from *Cellpose3* and the starting trained model used was the *Cyto2*. For this analysis, we used the *QuPath* extension for *Cellpose*, allowing us to streamline the workflow by combining *QuPath’s* visualization tools with the *Cellpose* segmentation model trained. Annotations created in *QuPath* were exported as a CSV file. The segmentation results were then used as input for further downstream analyses. Specifically, the segmented data were inputted into PCF and MERSCOPE platforms for spatial-transcriptomics and spatial-proteomics analysis, enabling the identification of spatial patterns and cell-to-cell interactions at single-cell resolution. We Inputted the segmentation mode in the Vizualizer tool (Vizgen), which provided an intuitive interface for examining the quality and accuracy of the segmentations. This visualization step was critical for ensuring that the segmentations aligned with biological expectations and tissue architecture.

### *AstroSuite for* Auto Assignment of Cell Identities and States, TCNs, and MCIMs

*AstroSuite* contains interconnected algorithms for analyzing spatial biology; two of the algorithms, *TACIT* and *Astrograph* have been published previously^26^. *<u>TACIT (Threshold-based Assignment of Cell Types from Multiplexed Imaging Data) for cell type annotation</u>*. We used *TACIT* (<u>T</u>hreshold-based <u>A</u>ssignment of <u>C</u>ell <u>T</u>ypes from Multiplexed Imaging Da<u>T</u>a) for cell type annotation in our spatial omics dataset. *TACIT* is an unsupervised algorithm designed for single-cell resolved spatial modalities, including spatial-transcriptomics and proteomics. *TACIT* takes as input the normalized CELLxFEATURE matrix as a result of *Cellpose3* cell segmentation and quantification of individual cell features such as probe intensity or count values. Additionally, a TYPExMARKER matrix, informed by expert knowledge, is used to indicate the relevance of specific markers for defining cell types. The annotation process occurs in two stages. In the first stage, cells are grouped into Microclusters (MCs) using the Louvain algorithm to capture highly homogeneous cell populations. Simultaneously, Cell Type Relevance scores (CTRs) are calculated for each cell through a linear combination of its normalized marker intensity vector with predefined cell type signatures. A higher CTR indicates a stronger association with a specific cell type and vice versa. Next, TACIT determines a threshold that distinguishes true positive cell type signals from background noise. CTRs are ranked, and segmental regression is applied to divide the CTR growth curve into 2 to 4 segments, identifying high and low relevance clusters. Unavoidably, there exists cells with low relevance score present in high relevance group and vice versa. An absolute CTR threshold is then determined to minimize such inconsistent assignments. Subsequently, cells with CTRs exceeding this threshold are labeled as positive for the associated cell type. In cases where cells are labeled with multiple cell types, *TACIT* employs a deconvolution step using the k-nearest neighbors (k-NN) algorithm and cells with unique cell type assignment to resolve such ambiguity. *<u>Constellation for Tissue Cellular Neighborhood Assignment in Whole Slide Imaging:</u>* We developed *Constellation* to address the challenges of unsupervised Tissue Cellular Neighborhood (TCN) assignment in Whole Slide Imaging (WSI) data, building upon the original CytoCommunity framework. While CytoCommunity effectively clusters cells within smaller tissue slides, it struggles with scalability for large WSIs or multiple slides. Constellation introduces novel partitioning strategies and a consensus-based assignment approach, enabling efficient processing of large datasets. The first step is WSI partitioning, where tissues are divided into smaller, non-overlapping sub-regions to reduce computational complexity. The tissue, represented as a set of cells 𝑊, with spatial coordinates and phenotypic labels, is partitioned using three strategies: vertical stripes (based on 𝑥-coordinate boundaries), horizontal stripes (based on 𝑦-coordinate boundaries), and square grids (dividing along both axes). Each partition contains a minimum and maximum number of cells to ensure computational efficiency and biological relevance, where partitions exceeding the maximum cell count are subdivided, and smaller ones are merged. Formally, let 𝑊 = ⋃^𝐾^_*i*=1_ 𝑆_𝑖_, where 𝑆_𝑖_ representing the 𝑖 − 𝑡ℎ subset of cells, and 𝐾 is the total number of partitions such that 𝑆_𝑖_ ∩ 𝑆_𝑗_ = ∅ for all 𝑖 ≠ 𝑗. Next, sub-graph 𝐺_𝑖_ = (𝑉_𝑖_, 𝐸_𝑖_) are constructed for each partition 𝑆_𝑖_ capturing both phenotypic and spatial relationships among the cells. Here, 𝑉_𝑖_ denotes the set of nodes (cells) within 𝑆_𝑖_, where each node 𝑣_𝑛_ ∈ 𝑉_𝑖_ is represented as a one-hot encoded phenotype vector:

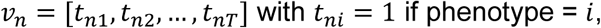

and 𝑇 is the total number of phenotypes. The edge set 𝐸_𝑖_ connects nodes based on the 𝑘-nearest neighbors determined by Euclidean distance in the spatial coordinate space. The sub-graph is represented by an adjacency matrix 𝐴_𝐺_*i*__ ∈ ℝ^|𝑉𝑖|×|𝑉_𝑗_|^, where entries are binary (1 for an edge, 0 otherwise), and a feature matrix 𝐹_𝐺_ ∈ ℝ^|𝑉𝑖|×𝑛^, representing cell phenotypes. The collective unsupervised training stage processes these sub-graphs in a graph neural network (GNN). Each sub-graph is treated as an independent training sample, and the model processes them in mini-batches with a gradient accumulation technique over 𝑁_𝑎_ = 32 batches. The GNN minimizes a combined loss function for training:

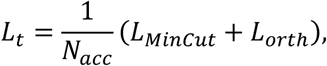

where *L_MinCut_* encourages separation between different clusters and 𝐿_𝑜𝑡ℎ_ promotes orthogonality, ensuring that each cell is confidently assigned to distinct TCNs. Finally, TCN assignment is performed using a consensus-based approach that aggregates results across the partitioning strategies. For each cell 𝑣_𝑛_, TCN assignments from vertical stripes, horizontal stripes, and square grid partitioning methods are compared, with the final assignment determined by majority voting:

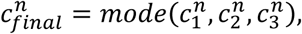

where *c^n^_m_* denotes the TCN assignment from the 𝑚-th partitioning method. In rare cases where all three assignments differ, the cell is excluded to maintain result reliability. This consensus-based approach ensures robust TCN assignments by integrating diverse spatial perspectives within the WSI. *<u>STARComm for spatial-resolved cell-cell communication:</u>* In STARComm (*<u>S</u>pa<u>TiA</u>l <u>R</u>eceptor-Ligand Analysis for Cell-Cell <u>COMM</u>unication)* analyses, we focus on identifying and characterizing co-location communication between cells based on 50 ligand-receptor (LR) interactions in spatial-transcriptomics datasets, such as MERSCOPE. The process begins by identifying cell pairs where communication may occur. Specifically, a cell that expresses a ligand gene (with value above 0 count) interacts with a nearby cell that expresses the corresponding receptor gene (also above 0 count), provided they are within 50 microns of each other in the spatial tissue layout. This process is repeated for all LR pairs (50 pairs in total), resulting in a network of interactions where each pair of cells that meets the ligand-receptor criteria is connected by an edge. Once all the LR interactions are established, we organize the spatial region into a grid of bins size 100 microns x 100 microns. Each bin in this grid captures the spatial region, and we ensure that each bin contains a very small number of edges representing ligand-receptor pairs. For each bin in the grid, we tally the count of individual LR pairs that fall within that spatial region, providing a cumulative measure of communication strength in that area. To analyze the distribution of these interactions more deeply, we calculate the kernel density for each ligand-receptor pair.

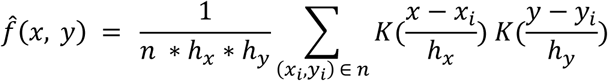

Where:

𝑓̂(𝑥, 𝑦): the density at the grid (𝑥, 𝑦)

𝑛: the total number of grids

ℎ_𝑥_ and ℎ_𝑦_ : the bandwidths in the 𝑥 and 𝑦 directions (default 10)

(𝑥_𝑖_, 𝑦_𝑖_) : the coordinates of the grid.

𝐾(.): kernel function - 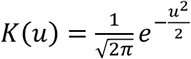

The kernel density estimation reveals how communication is spatially distributed—areas where the interactions between cells are more frequent will have higher density values, indicating that those locations are hubs of ligand-receptor communication. Finally, we apply the Louvain clustering algorithm to the grid, clustering the bins based on their communication density vectors. This clustering process groups spatial regions that exhibit similar co-location of LR pairs meaning that the LR interactions are concentrated in specific areas of the tissue. By clustering the bins, we capture and visualize how cell communication is spatially organized, highlighting areas where ligand-receptor interactions are occurring at higher densities, providing insight into the spatial dynamics of cellular communication across the tissue.

### *Astrograph* for spatial data visualization

We developed Astrograph, an R Shiny application that enables interactive visualization and analysis of spatial cell type data. The app allows users to customize cell type annotations with user-defined colors and select specific tissues or regions for focused visualization. It supports the generation of heatmaps that display cell type distributions based on marker expression, which can be tailored to reflect either cell type-specific markers or cell state markers. Beyond visualizing cell types, Astrograph also facilitates the exploration of cell states, providing insights into the proportions of various cell types and states across tissues or regions. Additionally, the app offers Voronoi plot functionality to depict spatial relationships between cells, thereby visualizing cell neighborhoods and interactions. Furthermore, Astrograph supports quantitative analysis of cell type interactions by counting neighboring relationships and identifying patterns in spatial cell dynamics, offering a comprehensive approach to spatial omics data exploration.

### Adapting *Drug2Cell*^70^ for spatial datasets

In the Spatial Drug2Cell analysis, we apply the Drug2Cell approach to the MERSCOPE data after the cells have been annotated with their respective cell types. Once cell type annotation is completed, we proceed by extracting the drug scores for each cell after we run drug2cell packages. These drug scores quantify the response or relevance of individual cells to specific drugs, based on expression profiles and other molecular characteristics. After obtaining the drug scores, we map these scores onto the spatial coordinates of the cells, allowing us to visualize the distribution of drug responses across the tissue landscape. This spatial mapping enables us to analyze how drug sensitivity or resistance varies based on the physical location of the cells within the tissue. Furthermore, we can compare the drug responses between different cell types and regions, highlighting variations in drug effectiveness or targeting based on cellular identity and spatial context. This comparison can reveal how certain cell types, such as immune cells, fibroblasts, or tumor cells, may exhibit distinct drug responses depending on their location within the tissue microenvironment. The combination of spatial information with drug scores provides a comprehensive view of how drugs interact with different cell populations, aiding in targeted therapeutic strategies.

### Motif graphs

The motif graph is a network-based representation designed to capture spatial adjacency relationships between groups of functional neighborhoods (FNs). A functional neighborhood can represent a specific TCN from Mega-CytoCommunity or an MCIM from StarComm. In this graph, each node corresponds to an FN, and an edge exists between two nodes if there exist neighboring cells linking two different FNs, i.e., a cell from one FN is within 50 micrometers of a cell in the other FN. Since FNs may appear in multiple locations on a slide as instances, the motif graph effectively summarizes the interaction relationships between these types of FNs, with repeated FN motifs forming a subgraph structure. For each pair of connected FNs we compute the proportion of neighboring cells linking them relative to the total number of cells in both FNs. This metric reflects the frequency of neighboring relationships, and it is visually represented by the thickness of the edges in the motif graph. Note that in the visualization of motif graph, we highlight only the top three most frequent interactions between clusters to emphasize key connections.

### GO and MSigDB Pathway Analysis

For all single-cell RNA-seq analyses, differentially expressed genes (DEGs) were identified using Trailmaker with a fold-change threshold of ≥0.5 and adjusted p-value <0.05 unless otherwise noted. DEGs were filtered for each fibroblast subtype or disease state comparison and subjected to Gene Ontology (GO) and MSigDB pathway enrichment using the Enrichr platform (https://maayanlab.cloud/Enrichr/). Only significantly enriched terms (adjusted p < 0.05) were retained for visualization and interpretation. For Figure 4 (niche-specific fibroblast programs) and Figure 6 (chronic periodontitis), analyses focused on identifying subtype-specific pathways related to ECM organization, immune surveillance, cytokine signaling, and stress response. For Figure 6, pathway enrichment was performed using a more stringent DEG threshold (fold-change ≥1.0, adjusted p-value <0.05) to reflect high-confidence spatial proteogenomic differences. Interactivenn (http://www.interactivenn.net/) was used to assess gene list overlap and visualize shared and unique gene sets between fibroblast subtypes across health and disease conditions.

## STATISTICAL METHODS

### General methods

A variety of tools and statistical approaches were used for data analysis, as appropriate for the data type and study objectives. Non-sequencing-based data were primarily analyzed using QuPath and/or Prism 9. The choice of statistical tests is detailed in the main text and figure legends. All statistical analyses were two-sided, and significance was determined at a threshold of (p-value<0.05). For comparisons between groups, a t-test was applied when normality assumptions were met. If the data did not meet these assumptions, a Wilcoxon rank-sum test (Mann-Whitney U test) was used instead. All visualizations, including graphs and supplementary figures, were created using Prism 9/10, a community instance of Trailmaker® (hosted by Parse), and the ‘ggplot2’, ‘pheatmap’, ‘ggforcè package in R unless explicitly stated otherwise. Venn Diagrams were generated using http://www.interactivenn.net. In spatial proteotranscriptomics analyses, workflows for cell segmentation, cell type annotation, and tissue structure annotation were tailored to the data and the biological questions of interest. Cell segmentation was performed using the advanced capabilities of Cellpose3, which excels at robust segmentation of complex tissue images. For annotation, AstroSuite was employed, providing an automated pipeline for assigning cell identities, functional states, tissue structure neighborhoods, and multiplexed cell interaction maps (MCIMs). These methods are described in detail in the “**Cell Segmentation for Multi-IF and MERFISH Images using *Cellpose3*** and ***AstroSuite for* Auto Assignment of Cell Identities and States, TCNs, and MCIMs**” sections. Heatmaps for spatial data were generated by summarizing marker expression. For PhenoCycler datasets, the mean expression levels were used to capture trends across the data, while for MERFISH datasets, the median count values were used to better handle the sparsity inherent in these datasets. When alternative methods for summarization were used, these are specified in the corresponding figure legends to ensure transparency and reproducibility. Differential expression analyses for genes, markers, and drug targets were conducted using the Seurat pipeline (version 5.1.0). This pipeline integrates preprocessing, normalization, and differential expression analysis, allowing us to compare expression profiles across groups effectively.

## DATA AVAILABILITY

All data, including links to original raw data from each of the 11 studies (oral mucosa^2,29–33^, salivary glands^32,34–36^, and pulp^33,37,38^) can be found at GEO: https://www.ncbi.nlm.nih.gov/geo/. Original raw data for soft palate, labial mucosa, and parotid salivary gland can be also found at GEO (TBD). The data can also be analyzed at: https://cellxgene.cziscience.com/collections/065ad318-59fd-4f8c-b4b1-66caa7665409

## CODE AVAILABILITY

Analysis notebooks and CELLxFEATURE matrices for MERSCOPE and Phenocycler-Fusion 2.0 data are available at: https://github.com/Loci-lab/Oral-Craniofacial-Atlas.

## ACKNOWLEDGEMENTS

IS and KMB want to extend a heartfelt thanks to the Human Cell Atlas Consortium, specifically Sarah Teichman, Aviv Regev, Ellen Todres, and Lucia Robson, as well as the HCA Oral & Craniofacial Bionetwork, for supporting this and ongoing project to map the oral and craniofacial tissues in health and diseases. We also want to thank OMAPiX, Inc, Psomagen and Histoserv for their thoughtful support of this and future projects. Furthermore, we acknowledge that this article has become stronger and more comprehensive through conversations with Katarzyna M Tyc (VCU), Di Wu, and Ji-Eun Park, both at UNC. Figure 1d, Supplementary Figure 1c, and Figure 4a were adapted using images from <u>BioRender.com</u> and released under a Creative Commons Attribution-NonCommercial-NoDerivs (CC-BY-NC-ND) 4.0 International license. Figure 1a, Supplementary Figure 1b utilized and adapted the “salivary-glands” icon by Servier (https://smart.servier.com/), which is licensed under CC-BY 3.0 (https://creativecommons.org/licenses/by/3.0/.) Data services in support of the research project were provided by the VCU Massey Comprehensive Cancer Center Bioinformatics Shared Resource. We would also like to thank the support and the staff of the NIDCR Office of the Clinical Director, NIDCR Dental Clinic, the National Eye Institute Ophthalmology Clinic, and the NCI Center for Cancer Research Anatomic Pathology. This work utilized the computational resources of the NIH HPC Biowulf cluster (http://hpc.nih.gov). We would like to thank the following NIH core facilities which enabled this study: NIDCR/NICHD Computational Biology and Genomics Core, the NIDCR Microscopy Core, the NIDCR Combined Technical Research Core, and the Laboratory of Genitourinary Cancer Pathogenesis (LCGP) Microscopy Core Facility. We would also like to thank the generous support of the NIH/UMD Graduate Partnership Program for TJFP. This research was principally supported through research awards to BMW from the Division of Intramural Research (DIR) Program of the National Institute of Dental and Craniofacial Research of the National Institutes of Health (NIH/NIDCR ZIA: DE000704). Massey and this work has been supported in part with funding from NIH-NCI Cancer Center Support Grant CA016059. DP is supported by Fundação para a Ciência e a Tecnologia (2020.08715.BD). IS funded by Barts Charity (MGU045), the Royal Society (RGS/R2/202291), and the British Skin Foundation (004/RA/23). This work was supported by generous start-up funds from the ADA Science & Research Institute (Volpe Research Scholar Award), Department of Oral and Molecular Craniofacial Biology, Philips Institute for Oral Health Research and VCU Massey Comprehensive Cancer Center (startup funds), and NIH/NIDCR 1RM1DE035338 to KMB. PRT was partly supported by the Chan Zuckerberg Initiative DAF award 2022-237918 and research grants from NIH/NHLBI (R01HL146557, R01HL160939, R01HL153375). We want to thank the Core Unit for Systems Medicine at the University of Würzburg (supported by the IZKF Würzburg, project Z-6). This work was supported by funding from the German Cancer Aid (MSNZ Würzburg/NG3) and the European Union/European Research Council (ERC Starting Grant number 101042738/OralNiche) to KK. This publication is part of the Human Cell Atlas: http://www.humancellatlas.org/publications.

## CONTRIBUTIONS

BFM, KLAH, DP, IS, JL, and KMB conceptualized the project.

BFM, KLAH, DP, AP, RS, PRT, XZ, MK, SAT, JL, KMB developed methods for data analysis.

BFM, DP, NK, QTE, BR, AG, SH, LS, MB, PP, ZK, NH, KIK, AK, SP, KK, BMW, IS, KMB supported sample collection.

BFM, KLAH, DP, XZ, MK, AVP, BTR, AP, RS, PRT, BN, MMM, IS, JL, KMB performed experimental analysis.

BFM, KMB wrote the original draft.

BFM, KLAH, DP, AP, XZ, QTE, SAT, RS, PRT, IS, JL, KMB reviewed and edited the final manuscript.

## KEY TERMS

AstroSuite: A modular collection of spatial bioinformatics tools (TACIT, Constellation, STARComm, and hist2omics) enabling integrated identification and visualization of cell types, features, and interactions across multimodal datasets.
Constellation: One of the components of the AstroSuite pipeline that defines tissue cellular neighborhoods (TCNs) by integrating all samples in an experiment and partitioning tissues into spatially co-occurring microenvironments enriched for coordinated cell activity.
hist2omics: One of the components of the AstroSuite pipeline for overlay and analysis of single-cell–resolved spatial omics data from the same slide with histology staining, here applied to spatial proteotranscriptomics combining spatial-proteomics, spatial-transcriptomics, and H&E.
Multicellular Interaction Modules (MCIMs): Spatial clusters co-localized receptor–ligand signaling; functional hubs analogous to TCNs but defined by active communication patterns.
Parainflammation: A low-grade, persistent activation of the immune system in tissues, positioned between homeostasis and classical inflammation.
Proteotranscriptomics: Spatial measurement of both protein and RNA expression from the same tissue sections at single-cell resolution.
Receptor–Ligand Density Scoring: A STARComm method quantifying spatial density and strength of receptor– ligand interactions in integrated tissue datasets.
STARComm: A computational framework quantifying receptor–ligand communication and identifying MCIMs based on communication density spatial expression density and proximity.
Structural Immunity: Immune regulation by structural cells (e.g., fibroblasts, endothelial cells) through spatial organization and signaling.
TACIT: An AI-enabled, automated whole-slide classification tool assigning cell types and tissue features from multiplexed spatial imaging data.
Tiered Cell Annotation: A multi-level cell labeling approach beginning with broad categories and refining to specific identities based on marker expression and tissue context.
Tissue Cellular Neighborhoods (TCNs): Recurring, spatially organized groups of co-occurring cell types predicted to coordinate local biological functions and signaling.

